# Leveraging IFNγ/CD38 regulation to unmask and target leukemia stem cells in acute myelogenous leukemia

**DOI:** 10.1101/2023.02.27.530273

**Authors:** Mariam Murtadha, Miso Park, Yinghui Zhu, Enrico Caserta, Ada Alice Dona, Mahmoud Singer, Hawa Vahed, Theophilus Tasndoh, Asaul Gonzalez, Kevin Ly, James F Sanchez, Arnab Chowdhury, Alex Pozhitkov, Lucy Ghoda, Ling Li, Bin Zhang, Amrita Krishnan, Guido Marcucci, John Williams, Flavia Pichiorri

## Abstract

Elimination of drug-resistant leukemia stem cells (LSCs) represents a major challenge to achieve a cure in acute myeloid leukemia (AML). Although AML blasts generally retain high levels of surface CD38 (CD38^pos^), the presence of CD34 and lack of CD38 expression (CD34^pos^CD38^neg^) are immunophenotypic features of both LSC-enriched AML blasts and normal hematopoietic stem cells (HSCs). We report that IFN-γ induces CD38 upregulation in LSC-enriched CD34^pos^CD38^neg^ AML blasts, but not in CD34^pos^CD38^neg^ HSCs. To leverage the IFN-γ mediated CD38 up-regulation in LSCs for clinical application, we created a compact, single-chain CD38-CD3-T cell engager (CD38-BIONIC) able to direct T cells against CD38^pos^ blasts. Activated CD4^pos^ and CD8^pos^ T cells not only kill AML blasts but also produce IFNγ, which leads to CD38 expression on CD34^pos^CD38^neg^ LSC-enriched blasts. These cells then become CD38-BIONIC targets. The net result is an immune-mediated killing of both CD38^neg^ and CD38^pos^ AML blasts, which culminates in LSC depletion.

**Statement of significance:** This work represents a potential advancement in the treatment of AML, as it involves the release of IFN-γ by T cells to induce CD38 expression and thus sensitizing leukemia stem cells, which have been resistant to current treatment regimens, to CD38-directed T cell engagers.

## Introduction

Despite recent FDA-approved new therapeutics, only 30% of adult patients with acute myeloid leukemia (AML) are alive at 5 years from diagnosis.^1,2^ Both treatment refractoriness and post-remission relapse remain the major challenges for these patients^3,4^. Allogeneic hematopoietic stem cell transplantation (alloHSCT) is the only curative approach for most of these patients, but treatment-related morbidity and mortality limit its use to clinically fit or younger patients ^5,6^. Thus, the development of more effective and less toxic therapies remains a major unmet need for AML patients.

In AML, the leukemia stem cell (LSC) population comprises primitive leukemic cells capable of self-renewal and disease initiation and maintenance.^7^ These cells are highly resistant to current AML therapeutics,^8^ and therefore they are responsible for treatment failures in AML patients. Compared with bulk AML blasts, LSCs are characterized by quiescence, distinct metabolism (i.e, they are highly dependent on oxidative phosphorylation), and phenotypic plasticity.^9^ LSCs are quite difficult to identify and target, given that these cells share common membrane antigens with non-stem leukemic cells (e.g., bulk blasts) and with normal hematopoietic stem cells (HSCs).^10^ Although they can reside in immunophenotypic-diverse leukemic cell subpopulations,^11^ LSCs are recognized to be enriched in the CD34^pos^CD38^neg^ fraction, while the CD34^pos^CD38^pos^ fraction mainly comprises more differentiated AML progenitors and bulk AML blasts.^9^

CD38 is an ectoenzyme transmembrane glycoprotein involved in the metabolism of extracellular nicotinamide adenine dinucleotide (NAD+).^12^ Loss of CD38 associates with poor immunogenicity and other metabolic changes, as CD38 activity reduces NAD_+_^13,14^ and affects cyclic ADP-ribose levels.^15^ CD38 has been exploited as a therapeutic target in multiple myeloma (MM) and T-cell acute lymphoblastic leukemia (T-ALL), where anti-CD38 monoclonal antibodies have demonstrated significant clinical activity,^16,17^ but in AML it remains to be fully explored.

Bispecific T cell engagers are antibodies that redirect host T cells to kill cancer cells by simultaneously binding a tumor associated antigen (TAA) on cancer cells and the CD3 receptor on T cells. Proofs of concept of this new class of drugs has recently emerged from the treatment of solid and hematological cancers, including MM and ALL.^18^ In AML, several TAAs including CD33, CD123, CD13, and CLL-1 have been shown to represent suitable targets for T cell engagers,^19^ but to date the feasibility of this therapeutic needs to be fully elucidated, given the immunophenotypic heterogeneity of AML cells and the overlap of these membrane proteins with those on normal HSCs ^20^.

Here we show that our newly generated single chain CD38 T cell engager (CD38-CD3 BIONIC) has a strong antileukemic activity against both CD38^pos^ and CD38^neg^ AML blasts. We found that this effect relies on T cell-dependent IFNγ degranulation, which coverts CD34^pos^CD38^neg^ into CD34^pos^CD38^pos^ blasts and results in leukemia eradication.

## Results

### CD38 is dynamically expressed and regulates stemness in AML

Aligned with previously published data,^21–23^ we observed that CD38 is expressed at significantly higher mRNA and protein levels in AML blasts compared to levels in normal counterpart cells (healthy donor, HD) (**Fig.1A,1B**), whereas a smaller fraction of the bulk leukemic population lacks surface expression of this antigen **(Fig.1C).** Interestingly, CD34^pos^CD38^neg^ AML blasts acquire CD38 expression (p = 0.008) when cultured *ex vivo* (**Supplementary Fig. S1A-S1C)** or engrafted in immunodeficient NSG mice (**Supplementary Fig.S1D-S1F**), thereby supporting the relevance of the surrounding microenvironment in regulating the levels of expression of this protein. Of note, CD38 negativity associates with higher “stemness” levels, as LSK (i.e., Lin^-^Sca^+^Kit^+^) cells from CD38 knockout mice (CD38^KO^),^24^ upon transduction with the AML-related onco-fusion gene *MLL-AF9^25^* (**Fig.1D**), have enhanced colony forming units (CFU) capacity *in vitro* **(Fig.1E,1F),** and exhibit higher bone marrow (BM) (p = 0.01) and spleen (p<0.0001) engraftment (**Fig. 1G–1J**) and splenomegaly (p = 0.0009) *in vivo* (**Fig. 1K**), as compared with levels from *MLL-AF9*-transduced CD38^pos^ LSK controls ^26^.

**Figure 1.**
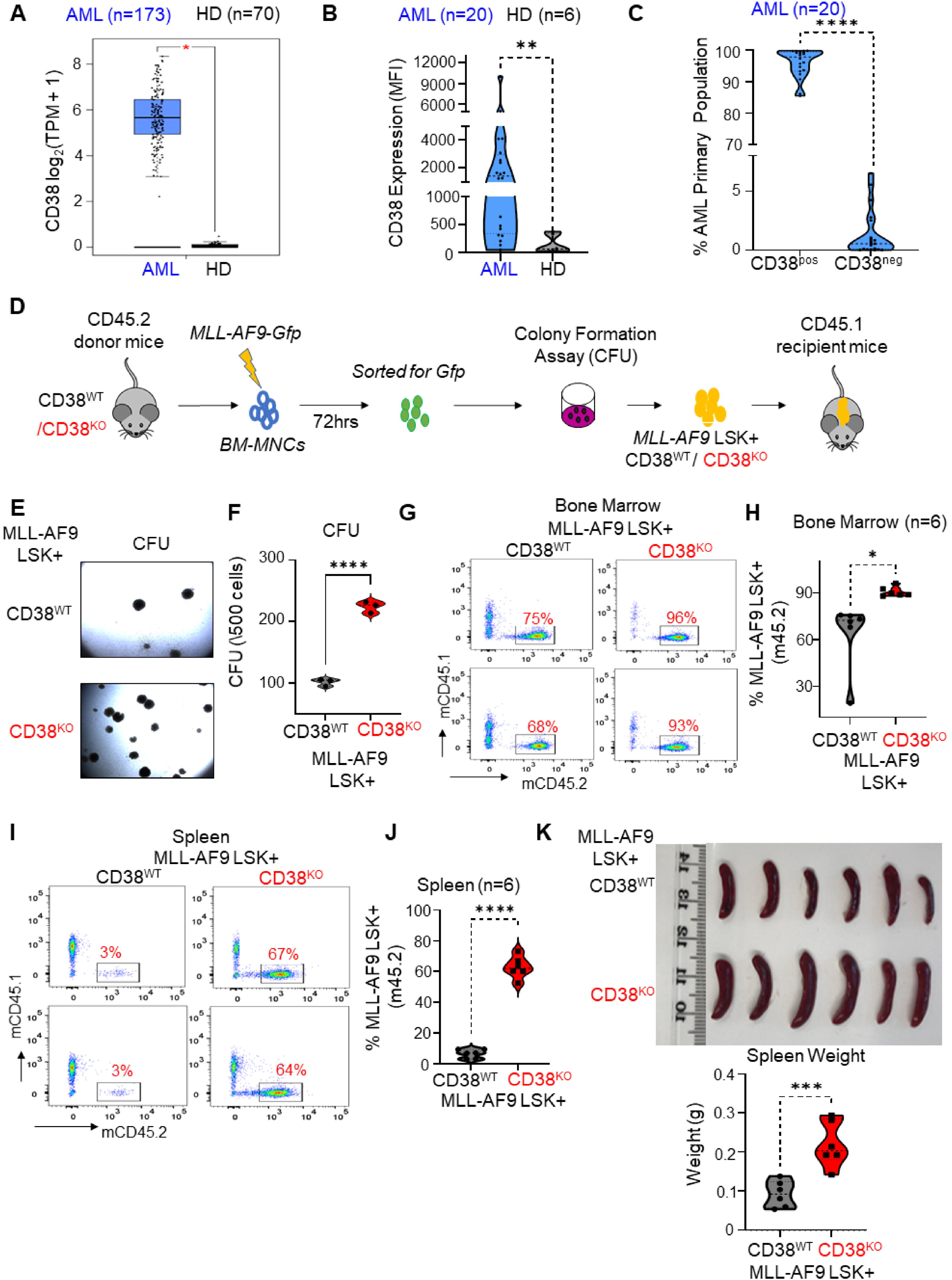
CD38 is dynamically regulated in AML. (A) CD38 mRNA expression in AML (blue, 173 samples) or HD (black, 70 samples) determined using gene expression profiling interactive analysis (GEPIA). Data for AML patients were downloaded from TCGA using LAML dataset; data for normal bone marrow were downloaded from GTex. (B) Violin plot comparing surface density of CD38 in total MNCs of AML patients (n = 20: 8 BM, 8 PB, and 4 LP samples) versus HD samples (n = 6 BM). Surface staining CD38-PE and CD45-APC was performed upon thawing of patient and HD MNCs. MFI is plotted for each patient and HD. Statistical significance was calculated using Mann-Whitney. **p<0.01. (C) Violin plot displaying percent of CD38^pos^ and CD38^neg^ cells in AML patients (n=20: 8 BM, 8 PB, and 4 LP samples). The percent of CD45^Dim^CD38^pos^ and CD45^Dim^CD38^neg^ cells were determined using three color CD34-FITC, CD38-PE, and CD45-APC surface staining followed by flow cytometry upon thawing of sample. Unpaired student t-test was used for statistical analysis. ****p<0.0001. (D) c-Kit^+^ cells were isolated from BM of CD38^WT^ and CD38^KO^ CD45.2 mice and transduced with *MLL-AF9-GFP* retroviral vector (MOI 5) for 72 hours. At 72 hours, GFP positive cells were sorted and plated for CFC assay. At day 10, leukemic colonies were harvested and transplanted into CD45.1 expressing congenic recipients to assess the role of CD38 in AML initiation. (E) Representative CFC assay images of CD38^WT^-MLL-AF9 and CD38^KO^-MLL-AF9 c-kit^+^ cells comparing their colony initiating capacity. (F) Violin plot comparing CFU capacity of CD38^WT^-MLL-AF9 and CD38-MLL-AF9 c-kit^+^ cells. Unpaired student t-test was used for statistical analysis. ****p<0.0001. (G-J) Representative flow cytometry dot plots and violin plots comparing engraftment of CD38^WT^-MLL-AF9 and CD38^KO^-MLL-AF9 c-kit^+^ cells in BM and spleen of recipient CD45.1 mice at harvesting (week 5 of transplantation). Unpaired student t-test was used for statistical analysis. *p<0.05; ****p<0.0001. (K) Spleen images showing prominent splenomegaly in CD38^KO^-MLL-AF9 CD45.1 recipient mouse. Violin plot comparing splenomegaly in CD38^WT^-MLL-AF9 and CD38^KO^-MLL-AF9 transduced mice. Unpaired student t-test was used for statistical analysis. ***p<0.001. (MNCs = mononuclear cells; HD = healthy donor; PB = peripheral blood; BM = bone marrow; LP = Leukapheresis; MFI = median fluorescence intensity; CFC = Colony forming cell assay; CFU = colony forming unit).

### IFN-γ induces CD38 expression in AML cells irrespective of their CD38 status

Recent reports have shown that IFNγ upregulates CD38 expression on CD38^pos^ AML cells ^27^ and blocks leukemia progression through type I innate lymphoid cells.^28^ To this end, we observed that *ex vivo* exposure of primary BM AML blasts to IFNγ resulted in a significant increase in CD38 protein levels in both CD34^pos^ and CD34^neg^ AML blasts (**Fig.2A–2C**) and reduction of cell clonogenicity (**Fig. 2D**) independently of their cytogenetics, including TP53 mutational status (**Supplementary Table S1**); these effects were significantly less evident in normal CD34^pos^CD38^neg^ BM HSCs (**Fig.2A–2D)**. Single cell RNA sequencing analysis of total BM mononuclear cells (BM-MNCs) isolated from either AML or healthy donors show that IFNγ treatment caused major transcriptional perturbation in the AML cells associated with significant transcriptional changes in more than 1000 genes (**Fig. 2E**, **Supplementary Fig. S2A**, **and Supplementary Table S2**), an effect that was not observed in the healthy immune subsets (**Fig. 2E and Supplementary Fig. S2B**). Specifically, we observed that IFNγ treatment induced significant changes in the oxidative phosphorylation and mitochondrial disfunction pathways in the AML cells, but these changes were not found in any of the normal immune subsets analyzed (p<0.0001, **Supplementary Fig. S2C and Supplementary Tables S3 and S4**). CD38 mRNA expression was also significantly upregulated in AML cells upon exposure to IFNγ (**Supplementary Table S2**). To validate these results, we then performed bulk RNA sequencing in the CD34^pos^ cells isolated from the BM-MNCs of relapsed AML patients (n=3) and healthy donors (n=3) treated overnight with or without 10 ng/ml of IFNγ. Hierarchical clustering analysis showed that IFNγ treatment induced a significant upregulation of antigen processing and presentation related genes in both AML and healthy CD34^pos^ cells compared to the untreated counterparts (**Fig. 2F**). Gene set enrichment analysis (GSEA) also shows induction of IFNγ signaling related pathways in both AML patients and healthy donors (**Fig. 2G**). We additionally observed that, upon IFNγ treatment, CD34^pos^AML cells, but not HD cells, display the upregulation of genes involved in the oxidative phosphorylation (**Fig. 2H**), fatty acid metabolism, adipogenesis, MTOR signaling, peroxisome, and adipogenesis pathways (**Supplementary Fig. S2D**) and the downregulation of genes involved in AML cell survival such as CD109^29^, KDM6B,^30^ and MYCN^31^ (**Supplementary Fig. S2E)**, which are normally upregulated in AML compared to normal cells. Higher CD38 mRNA expression was observed in CD34^pos^AML cells compared to the healthy counterpart (**Supplementary Fig. S2F**).

**Figure 2.**
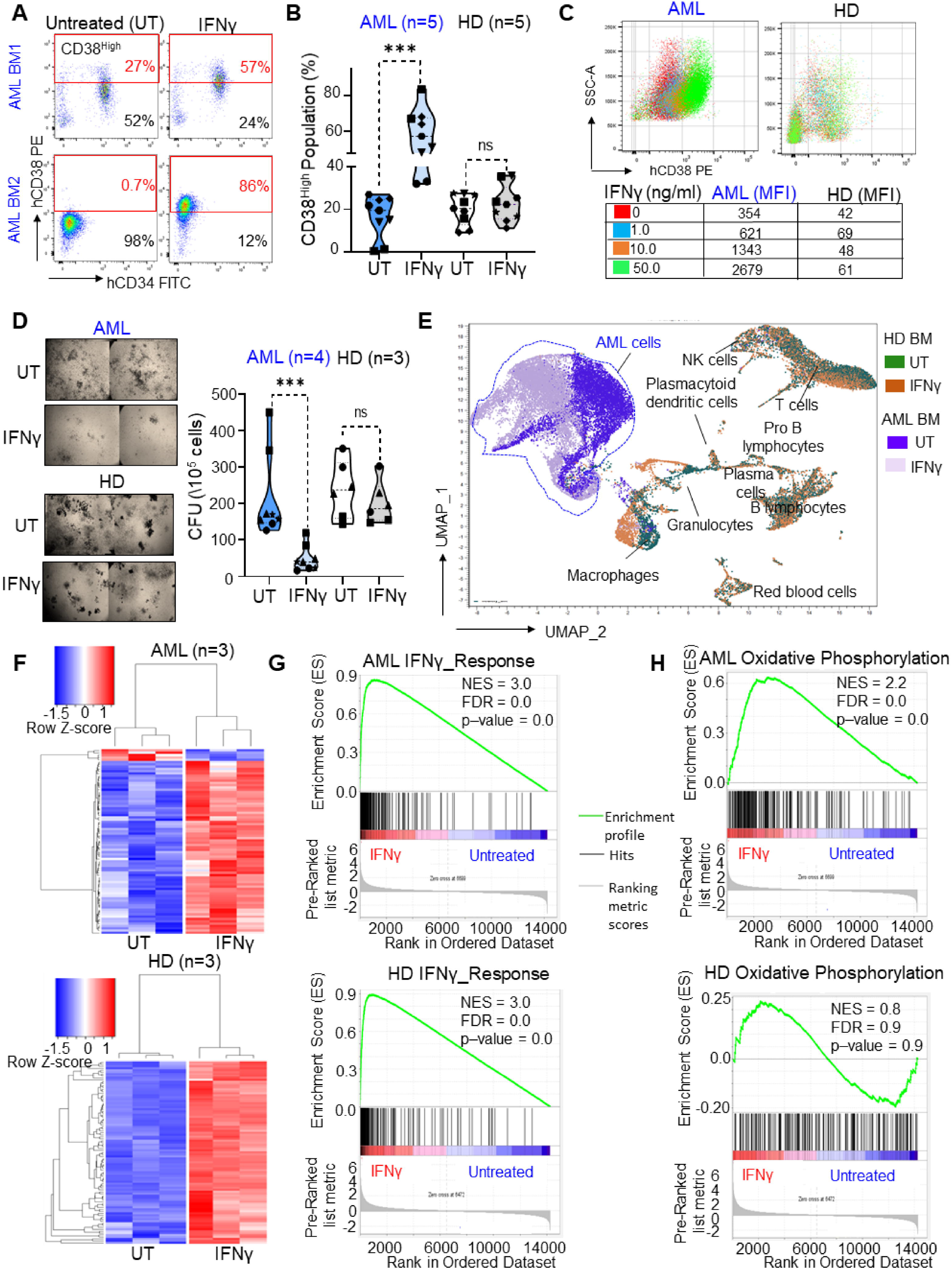
IFN-γ induces CD38 expression in AML cells irrespective of their CD38 status. (A) Representative flow cytometry dot plots for 2 AML BM samples comparing percent of CD38^High^ population in IFNγ treated group versus untreated. IFNγ was added overnight at a concentration of 10 ng/ml. (B) Violin plot comparing percent of CD38^High^ population between untreated and IFNγ treated groups for AML patients (n=5: 4 BM and 1 PB) and HD (n = 5 BM). IFNγ was added overnight at a concentration of 10 ng/ml. Each shape represents a different AML patient and HD. Each patient and HD sample were done in duplicates. Paired student t-test was used to calculate statistical significance between untreated and IFNγ treated groups for AML patients and HD. ***p<0.001; ns, not significant. (C) AML and HD BM MNCs were treated overnight with different doses of IFNγ (1.0, 10.0, and 50.0 ng/ml), and CD38 expression was determined in CD45^Dim^ population with flow cytometry. Overlaid dot plots show shift in CD38 expression with IFNγ treatment, and the table below lists MFI for each dose and control. (D) Representative images of CFC assay for AML patient and HD BM MNCs following treatment with IFNγ (10 ng/ml). Images were acquired at 2X magnification to cover complete well and stitched. Violin plot comparing colony forming units (CFU) between untreated and IFNγ (10 ng/ml) treated groups for AML patients (n= 4: 3 PB and 1 BM) and HD (n= 3 BM). Each shape represents different AML patient and HD. Each patient and HD were done in duplicates. Paired student t-test was used to calculate statistical significance between untreated and IFNγ treated groups for AML patients and HD. ***p<0.001; ns, not significant. (E) UMAP representation combining one AML patient and one HD BM-MNCs to depict changes in immune cell subsets and AML cells after IFNγ treatment for 5 hours at 10 ng/ml. (F-H) CD34^pos^ cells were purified from 3 AML patients and 3 HD BM samples, and cells were plated onto two wells for each patient and HD, where one well was untreated and other was treated with IFNγ at 10 ng/ml. The next day, cells were collected, and RNA was extracted and subjected to bulk RNA-seq. (F) Z-score based hierarchical clustering heatmap showing top 75 upregulated genes and 5 downregulated genes in 3 AML patients upon IFNγ treatment and differentially upregulated 93 genes in HD cells (n=3) upon IFNγ treatment (p<0.05). (G) GSEA generated enrichment plots of IFNγ signaling pathway that was activated in HD and AML CD34+ cells. (H) GSEA generated enrichment plots of oxidative phosphorylation pathway that was exclusively upregulated upon IFNγ treatment in AML patients and not HD. The enrichment score (ES) is calculated based on the pre-ranked gene list obtained by “-log10(*p* value for DEG)□×□sign(log2(fold change))” (FDR<0.05). (MNCs = mononuclear cells; HD = healthy donor; PB = peripheral blood; BM = bone marrow; MFI = median fluorescence intensity; CFC = Colony forming cell assay; CFU = colony forming unit).

### Generation of a novel CD38 T-cell engager (CD38-CD3 BIONIC)

Because CD4^pos^ T helper type 1 and CD8^pos^ cytotoxic T cells release IFN-γ^32^ when they engage their target(s) and become activated, we hypothesized that activation of T cells with a CD38-directed T cell engager could not only directly kill CD34^pos^CD38^pos^ AML blasts but also induce high levels of IFN-γ secretion and in turn convert the CD34^pos^CD38^neg^ LSC-enriched fraction into CD34^pos^CD38^pos^ blasts, which would themselves become a direct target of the T-cell engager (**Fig.3A**). The expected net result could be not only significant cytoreduction of leukemic blasts, but also depletion of LSCs that reside in the CD34^pos^CD38^neg^ fraction.

**Figure 3.**
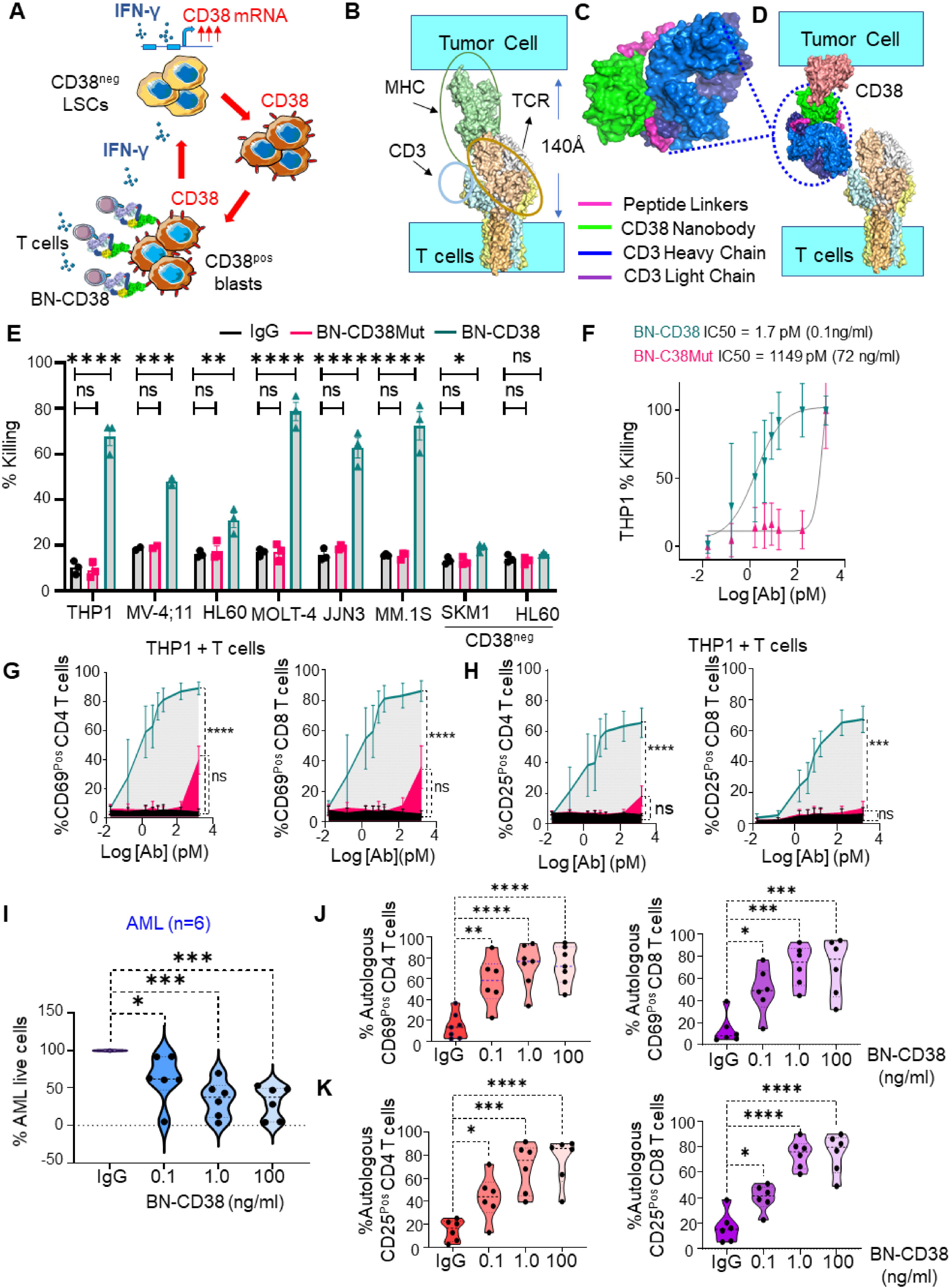
Generation of a novel CD38 T-cell engager (CD38-CD3 BIONIC). (A) Graphical representation of the therapeutic rationale to target CD38 LSCs with T cell engagers against CD38 to eradicate AML by uncovering the IFNγ/CD38 regulatory loop. (B) Graphical representation depicting distance between T cell and tumor cell membranes upon TCR-MHC interaction. (C) Scheme of BN-CD38 and projected structure with heavy chain of anti-CD3 Fab (blue), CD38 nanobody (bright green), light chain of anti-CD3 Fab (purple) and linkers (magenta). (D) Graphical representation of BN-CD38 interaction with T cell (CD3) and tumor cell (CD38). Predicted spatial orientation of BN-CD38 is similar to MHC-TCR complex. (E) CD38+ THP1, MV-4;11, MOLT-4, MM.1S, and CD38^neg^ SKM1 (GFP^pos^ or CFSE labelled) were co-cultured with healthy donor T cells at an E:T ratio of 1:1 overnight (16hrs) in the presence of 1.0 ng/ml of BN-CD38, BN-CD38Mut, and control human IgG. HL60 and HL60 CD38 CRISPR knockout cell lines were treated with 1 μM ATRA for 48 hours before starting co-culture, as is necessary to induce CD38 expression in HL60 cells. Cancer cell killing was assessed with 7-AAD staining and gating of GFP^pos^ or CFSE^pos^ cells. Bar graphs are represented as mean ± SEM of 3 healthy donors. Ordinary one-way ANOVA with Dunnett’s multiple comparisons test was used to calculate statistical significance. *p<0.05; **p<0.01 ***p<0.001; ****p<0.0001; ns, not significant. (F) THP1 GFP^pos^ cells were co-cultured with healthy donor T cells at an E:T ratio of 1:1 overnight (16hrs) in the presence of increasing doses of BN-CD38 and BN-CD38Mut. THP1 cell killing was determined with 7-AAD staining and gating on GFP^pos^ cells by flow cytometry. IC_50_ curves are shown for BN-CD38 and BN-CD38Mut, and data are represented as mean ± SEM of 5 independent healthy donors. (G-H) THP1 GFP^pos^ and healthy donor T cells were co-cultured at an E:T ratio of 1:1 overnight (16hrs) in the presence of different doses of BN-CD38, BN-CD38Mut, and control human IgG antibodies. Early and late T cell activation was assessed by surface expression of CD69 and CD25, respectively, in CD4^pos^ and CD8^pos^ subsets with flow cytometry and by excluding GFP^pos^ cancer cells. Dose-dependent T-cell activation curves are represented as mean ± SEM of 3 independent healthy donors. Ordinary one-way ANOVA with Dunnett’s multiple comparisons test was used for calculation of statistical significance. ***p<0.001; ****p<0.0001; ns, not significant. (I) Violin plot showing dose-dependent decrease of AML cells upon redirecting autologous T cells with BN-CD38 compared to IgG control. AML patient MNCs (n= 6: 4 PB and 2 BM, 3 newly diagnosed and 3 relapsed patients) were treated with BN-CD38 and control human IgG for 5 days. %CD45^Dim^CD34^pos^ or %CD45^Dim^CD33^pos^ was normalized to control human IgG for each patient. Ordinary oneway ANOVA with Dunnett’s multiple comparisons test was used to calculate statistical significance. *p<0.05; ***p<0.001. (J-K) Violin plots showing dose-dependent early (CD69) and late (CD25) T cell activation in autologous CD4^pos^ and CD8^pos^ T cells from AML patients, when treating with BN-CD38, in contrast to that from control human IgG. AML patient bulk MNCs (n= 6: 4 PB and 2 BM) were treated with increasing doses of BN-CD38 and 100 ng/ml of control human IgG for 48 hours and subjected to CD4, CD8, CD69, and CD25 surface staining followed by flow cytometry analysis. Ordinary oneway ANOVA was used to calculate statistical significance. *p<0.05; **p<0.01; ***p<0.001; ****p<0.0001. (MNCs = mononuclear cells; HD = healthy donor; PB = peripheral blood; BM = bone marrow; Target cancer cells, T; Effector T cells, E).

To this end, we created a highly compact and efficient antileukemic CD38-CD3 T cell engager. Superimposing the cryo-electron microscope structure of the T cell receptor (TCR) (PDB: 6JXR) and the major histocompatibility complex (MHC) structure (PDB:4GRL), we first calculated that the distance between T cell membrane and antigen presenting cells (APC) is ~140 Å (**Fig. 3B**). To closely match the distance between the TCR and CD38 to that of the innate TCR/MHC interaction, the structure of which allowing for an efficient engagement of both CD8^pos^ and CD4^pos^ T cells, we inserted a CD38 nanobody between the light and heavy chains of an anti-CD3 Fab using two short peptide linkers. The net result of this design process is a compact, single-chain molecule we called CD38-CD3 BIONIC (<u>bio</u>logics <u>n</u>ested <u>i</u>nside <u>c</u>hains, BN-CD38) (**Fig.3C and Supplementary Fig. S3A)**, which also allowed us to bypass the complicated assembly issues inherent with the chain pairing of bispecific IgG. Through simple modeling, we showed that BN-CD38 indeed closely matched the TCR-MHC distance (**Fig.3D**). Surface plasmon resonance (SPR) and analytical cytometry studies showed that BN-CD38 retains binding affinity to both CD38 and CD3 antigens (**Supplementary Fig. S3B and S3C**). As a control, we also generated a BN-CD38 mutant (BN-CD38Mut) containing a single point mutation replacing serine with histidine at residue 57 in the binding pocket of the anti-CD38 nanobody (**Supplementary Fig. S3D**). BN-CD38Mut bound to CD38 significantly less than BN-CD38 (**Supplementary Fig. S3E**), while binding to CD3 remained unaffected (**Supplementary Fig. S3F**). When T cells (effector, E) were treated with BN-CD38 and co-cultured with CD38^pos^ (target, T) cancer cells including AML (i.e., THP-1, MV-4;11, HL-60), MM (i.e., MM.1S, JJN3) and T-ALL (i.e., MOLT-4) (target, T) at an E:T ratio of 1:1, we observed a strong T cell killing activity (p<0.0001) in all but the CD38^neg^ cells (i.e., SKM1, an AML cell line) and in CD38^ko^ HL-60 AML cells, in which the CD38 encoding gene was silenced via CRISPR (**Fig. 3E and Supplementary Fig. S3G**). No T cell killing was observed when the CD38^pos^ cell lines were incubated with BN-CD38Mut or non-specific human IgG (**Fig. 3E**). The IC_50_ for BN-CD38 was calculated to be within the 0.01 to 0.1 ng/ml range, while BN-CD38Mut had an IC_50_ at least 100-fold higher (**Fig. 3F and Supplementary Fig. S3H and S3I)**. Significant induction of early (CD69) and late (CD25) T cell activation markers on both CD4^pos^ and CD8^pos^ T cells was observed only in the presence of BN-CD38 (p<0.0001), and not with BN-CD38Mut or IgG control (**Fig.3G,3H and Supplementary Fig. S4A andS4B)**. Of note, a significant increase in cytokine levels including IFNγ, TNFα, and IL-2, among others, was observed only when BN-CD38, but not BN-CD38Mut or IgG, treated T cells were co-culture with CD38^pos^ THP-1 cells (**Supplementary Fig.S4C and S4D**). Significantly lower cytokine release was also observed when BN-CD38 treated T cells were incubated with CD38^KO^-HL-60 cells, compared to the CD38^pos^ parental cell line (**Supplementary Fig. S4E**). The compact structure of BN-CD38 for engagement and activation of both CD4^pos^ and CD8^pos^ T cells resulted in comparable and even enhanced killing capabilities of CD4^pos^ T cells compared with cytotoxic CD8^pos^ T cells (**Supplementary Fig. S4F**).^33^ BN-CD38 also induced significantly higher levels of IFNγ in both CD4^pos^ and CD8^pos^ T cells (p<0.0001) compared to BN-CD38Mut and control IgG (**Supplementary Fig.S4G and S4H**). Dosedependent autologous T cell killing and activation from BN-CD38 were also observed in co-cultures of matched PB and BM-MNCs from newly diagnosed or relapsed AML patients (**Fig. 3I–3K, Supplementary Table S1**). BN-CD38 also significantly reduced clonogenicity of primary AML cells compared with control IgG (p=0.0007) but did not affect that of normal HSCs (**Supplementary Fig. 5A**).

### BN-CD38 eradicates leukemia *ex vivo* and *in vivo*

Next, to investigate the effect of BN-CD38 on the cellular composition of the BM samples from AML patients, we assembled a 36-metal mass cytometry (CyTOF) panel designed to distinguish subpopulations of hematopoietic and immune cells (**Supplementary Table S5 and S6**). While BM samples from AML patients was lymphodepleted compared with those from healthy donors (**Fig. 4A and Supplementary Fig. S5B**), using supervised gating on vi_SNE (FlowSOM) and data-driven self-stratifying clustering CITRUS analysis, we showed that treatment of AML BM samples with BN-CD38 for 5 days significantly reduced the total AML population (**Supplementary Fig. S5C**) including CD34^pos^CD38^pos^ and CD34^pos^CD38^neg^ AML blasts (**Fig.4A-C**); this effect was not observed in normal BM samples (**Supplementary Fig.S6A**). Using FlowSOM analysis, we also showed that BN-CD38 significantly expanded CD8^pos^ effector memory (EM) (p=0.026) and terminally differentiated effector memory CD45RA^pos^ cells (TEMRA) (p = 0.03) (**Fig. 4D**). A significant expansion of IFNγ-secreting memory Tregs (p=0.002), and NKT cells (p=0.015) (**Fig.4E and Supplementary Fig. S6B**) was also observed, further corroborating the ability of BN-CD38 to activate CD4^pos^ T cells against autologous AML blasts and induce IFN-γ secretion (p=0.04) (**Supplementary Fig. S6C**). Furthermore, BN-CD38, but not BN-CD38Mut or IgG control, induced a significant increase in levels of CD38 surface protein (p=0.02) and mRNA (p=0.002) in AML blasts regardless of their initial CD38 status, but not in healthy BM cells (**Supplementary Fig.S6D and S6E)**. To support the relevant antileukemic role of IFNγ from BN-CD38 activated T-cells, we then treated total MNCs isolated from a relapsed AML patient for five days with 1 ng/ml BN-CD38 and 2 μg/ml anti-human-IFNγ neutralizing antibody. Using CyTOF analysis, we showed that blocking IFNγ attenuated the antileukemic BN-CD38 activity and suppressed T cell expansion (**Fig.4F and 4G**). Consistent with these results, T cells engaged with BN-CD38 upregulated CD38 on the surface of THP1 cells; this effect was reversed by the IFNγ blocking antibody (**Supplementary Fig.S6F and S6G**). Next, we investigated the antileukemic activity of BN-CD38 *in vivo*. Accordingly, 10^6^ luciferase-positive (Luc+) CD38^pos^ THP1 cells were injected intravenously (IV) into NOD-*scid* IL2Rg^null^ (NSG) mice (**Fig. 4H**). On day 19 post-injection, mice with similar leukemia burden as measured by bioluminescence (BLI) were randomized into two groups and treated with either 2.5 mg/kg BN-CD38 (n=8) or human IgG (n=10), and 3×10^6^ human T cells were given IV weekly for 4 weeks. A significant decrease in leukemic burden as measured by BLI was observed in the BN-CD38-treated mice compared with human IgG-treated controls as early as one week after start of the treatment (p=0.003) **(Fig. 4I and Supplementary Fig. S7A).** While lymphoproliferation and graft-vs-host disease (GVHD) were relatively common side-effect of T-cell infusion, 38% of BN-CD38 treated mice remained AML- and GVHD-free up to 120 days after the last treatment, with a not-reached median survival **(Fig.4J)**. In contrast, all the control mice died by day 30. The specific anti-leukemic activity of BN-CD38 was also observed in mice xenografted with U937 cells, which have significantly lower CD38 expression (CD38^dim^) compared to THP-1 **(Supplementary Fig. S7B)**. The (Luc+) U937-engrafted mice treated with BN-CD38 and T cells, and not those treated with BN-CD38Mut and T cells, had a near complete disease eradication (**Supplementary Fig. S7C-S7F**), supporting BN-CD38 induced T cell-mediated leukemia killing despite the lower membrane density of the target antigen (i.e., CD38) on the leukemia cells.

**Figure 4.**
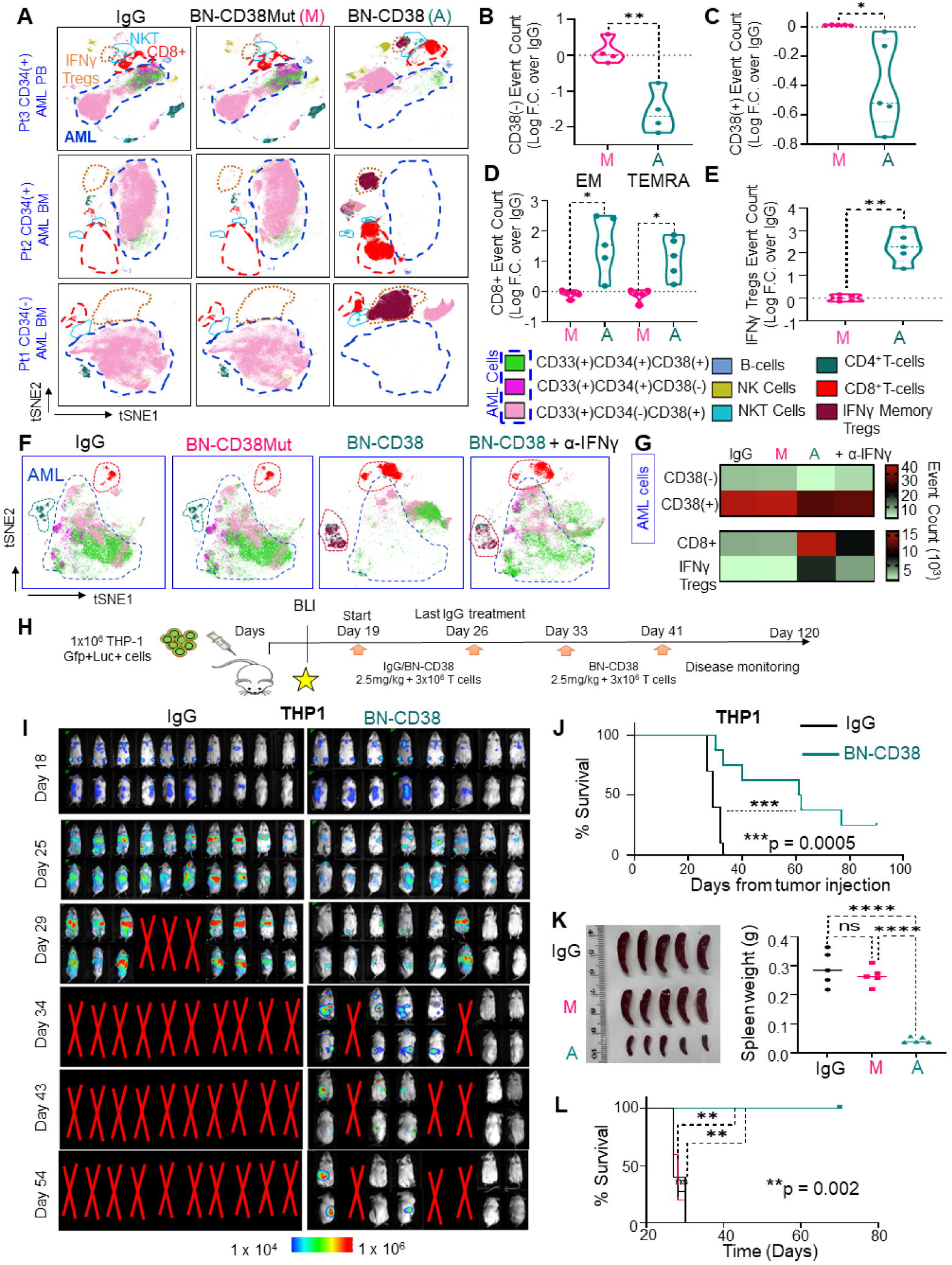
BN-CD38 eradicates leukemia *ex vivo* and *in vivo*. (A-E) Total MNCs from 5 AML patients (3 PB and 2 BM; 4 CD34^pos^ patients and 1 CD34^neg^ patient) were treated with 1.0 ng/ml of BN-CD38, BN-CD38Mut, control human IgG for 5 days and subjected to CYTOF immunophenotyping comprising 36 surface markers tailored to detect AML primary cells and different immune subsets. Analysis was performed with Cytobank© platform. (A) Supervised high-fidelity FlowSOM (“self-organizing maps”) based on vi-SNE 2D analysis for MNCs from 3 AML patients (1 PB and 2 BM) showing BN-CD38 reduces CD38^pos^ and CD38^neg^ AML cells and expands T cell subsets. FlowSOM parameters are: metacluster = 10 and cluster = 100, and viSNE parameters are: iterations = 1500-600 and perplexity = 50-100. Equal number of events were analyzed for each treatment group for each patient, and bulk MNCs were gated. (B-C) Violin plots showing BN-CD38, but not BN-CD38Mut, significantly decreased CD38^pos^ and CD38^neg^ AML cells event count. Event counts of BN-CD38 and BN-CD38Mut were normalized to control human IgG, and normalization was converted to log scale. (D-E) Violin plots showing BN-CD38, but not BN-CD38Mut, expanded CD8^pos^ effector memory (EM), terminally differentiated effector memory CD45RA^pos^ cells (TEMRA), and IFNγ-secreting memory Tregs. Event counts of BN-CD38 and BN-CD38Mut were normalized to control human IgG, and normalization was converted to log scale. (F-G) AML patient bulk MNCs were treated with 1.0 ng/ml of BN-CD38, BN-CD38Mut, control human IgG, and combination of 1.0 ng/ml BN-CD38 + 2 μg/ml αIFNγ for 5 days and subjected to custom 36-surface marker immunophenotyping by mass cytometry. (F) FlowSOM map of different AML and lymphocyte clusters for each treatment group. (G) Heatmap of event counts of AML and lymphocyte cell populations for each treatment group generated by FlowSOM. (H) One million THP1 cells stably expressing luciferase were intravenously (IV) injected into NSG mice. On day 18, engraftment was confirmed with bioluminescence imaging (BLI), and mice were randomized into two groups with comparable tumor burden (n=10 control human IgG and n=8 BN-CD38WT). Treatment was started on day 19, and mice were administered 2.5 mg/kg/mouse control human IgG or BN-CD38, together with 3 million healthy donor derived human T cells/mouse weekly by IV. A total of 4 treatments with BN-CD38 and two treatments with human control IgG were performed Mice were monitored weekly by BLI for assessment of tumor burden. (I) BLI data showing that BN-CD38 treated group had lower tumor burden and that 38% of the BN-CD38 treated group were cured. (J) Kaplan-Meier survival curve analysis showed BN-CD38 prolonged survival of mice. Logrank (Mantel-Cox) test was used to determine statistical significance. ***p=0.0005. (K-L) One million AML patient MNCs (Patient 14, Supplementary Table S1 were IV injected into each irradiated mouse. On day 20, engraftment of human cells was monitored, and mice were randomized into three groups, with each group including mice of comparable tumor burden: control human IgG (n = 5), BN-CD38 (n = 5), and BN-CD38Mut (n = 5). Treatment was started on day 21, and mice were administered 2.5 mg/kg/mouse control human IgG, BN-CD38, or BN-CD38Mut, together with 3 million healthy donor derived human T cells, which were given by IV weekly for 3 weeks. On day 42, BM and SP MNCs were harvested, and engraftment of human cells was assessed by flow cytometry. One million BM MNCs collected from each treatment group were transplanted in secondary mice by IV. Each week, mice were treated with 2.5 mg/kg/mouse of control human IgG, BN-CD38, or BN-CD38 Mut, together with 3 million healthy donor derived T cells by IV; a total of three treatments were given. On day 27, BM and SP MNCs were harvested, and engraftment of human cells was assessed. One million BM MNCs from each treatment group were IV transplanted in tertiary mice, and survival was monitored (PDX timeline is also shown in Suppl. Fig. 7G). (K) Images and scatter plot showing BN-CD38 suppressed splenomegaly in the PDX model on secondary BMT. Ordinary one-way ANOVA with Dunnett’s multiple comparisons test was used to calculate significance. ****p<0.0001; ns not significant. (L) Kaplan-Meier survival curve analysis showed BN-CD38 prolonged survival of PDX model on third BMT. Log-rank (Mantel-Cox) test was used to determine statistical significance. **p=0.002. (A = active treatment [BN-CD38], M = mutant [BN-CD38Mut], MNCs = mononuclear cells; PB = peripheral blood; BM = bone marrow; LP = leukapheresis; BMT = Bone marrow transplantation; PDX = patient derived xenograft; SP = spleen).

We also validated these results in patient derived xenograft models (PDXs). One million BM MNCs from two relapsed AML patient with complex karyotype (**Supplementary Table S1**) were injected into irradiated NSG mice (**Supplementary Fig.S7G**). On day 20, upon confirmation of successful engraftment by flow cytometry analysis, mice were randomized into three groups and treated weekly with BN-CD38, BN-CD38Mut or IgG (2.5 mg/kg/weekly) in combination with human T cells (3×10^6^/weekly) for a total of three weeks. To eliminate the confounding GVHD effect that would also be lethal to the disease-free mice, we sacrificed the mice after three treatments. The BN-CD38 treated mice had significantly reduced AML engraftment in the BM and spleen compared with that from BN-CD38Mut- or human IgG-treated controls (p<0.0001) (**Supplementary Fig. S7H**). We then proceeded with transplanting BM-MNCs of each of the treated animal groups into secondary NSG recipients. After 10 days, the transplanted mice received the same weekly treatment of the respective matched donors for three weeks. At the end of treatment, we observed that BN-CD38 treated groups had complete eradication of leukemia cells, both in the BM and in the spleen, compared to mice treated with BN-CD38Mut or IgG (p<0.0001) (**Fig. 4K and Supplementary Fig. S7I**). We then proceeded with a tertiary transplant experiment. We showed that mice engrafted with BM-MNCs from the BN-CD38-treated donors had no sign of leukemia as measured by flow cytometry analysis. Median survival was not reached, and 100% of the mice remained leukemia free; by contrast, transplanted mice treated with BN-CD38Mut or IgG had a median survival of 28 days each (**Fig.4L**). A significant reduction in engraftment was also observed after only two weeks of BN-CD38 treatment in mice transplanted with primary AML cells carrying complex karyotype and TP53 mutation, compared to effects with BN-CD38Mut or control IgG (**Supplementary Fig. S7J and S7K)**.

## Discussion

Unlike its role in MM and chronic lymphocytic leukemia (CLL), the role of CD38 in myeloid leukemogenesis remains to be fully elucidated. An earlier study including 304 newly diagnosed and relapsed AML patients showed that increased CD38 surface expression is associated with improved AML survival and favorable prognosis ^22^. Also, it was shown that the differentiating agent all trans retinoic acid (ATRA) induced CD38 expression in AML cell lines and primary patients, which implied CD38 is involved in AML cell differentiation ^34^.

In spite of the fact that CD38^neg^ blasts constitute a minor population in the total AML blasts,^23^ our data show that c-kit+ stem cells from CD38 knockout mice transduced with the AML-related onco-fusion gene *MLL-AF9* display enhanced leukemia cell engraftment, complementing previous studies demonstrating that lack of CD38 expression on AML cells is associated with enhanced leukemia initiating capabilities.^26^ We show that IFNγ induces CD38 mRNA and protein upregulation on AML cells independently of their initial CD38 expression. Notably, IFNγ also induces CD38 upregulation in AML blasts with complex karyotype including TP53 mutation, a genetic aberration that is associated with an extremely poor prognosis, specifically a median overall survival (OS) of 4 to 6 months and a 2-year OS rate <10% ^35^. Although previous studies have shown that interferon IFN type I (alpha and beta) and type II (gamma) induce CD38 transcriptional and protein surface expression on CD38^pos^ AML blasts and leukemia B cells independent of interferon regulatory factor-1 (IRF1),^36,37^ the exclusive upregulation of CD38 on AML cells, including CD34^neg^CD38^neg^ blasts but not in HSCs, has not been reported to the best of our knowledge.

Our results show that IFNγ-treated AML cells, and not normal HSCs, gained CD38 expression, which was associated with reduced clonogenicity. These data are also fully aligned with recently published results showing that IFNγ blocks leukemia progression through type I innate lymphoid cells in preclinical AML models.^28^ Gene expression profiling highlights that, in AML cells but not in HSCs, IFNγ induces the upregulation of metabolic pathways, including oxidative phosphorylation and fatty acid metabolism, which are critical to leukemia cell mitochondrial metabolism and survival.^38^ These results further highlight a metabolic difference of LSC cells versus HSCs, as also recently published.^38^ CD38 involvement in metabolic reprogramming in both cancer cells and immune effector cells through its NADase activity has been thoroughly reported.^13,39–41^ Hence, it is not surprising to find its selective surface upregulation upon IFNγ treatment in cells with altered metabolic pathways such as LSCs and blasts, in contrast to HSCs cells. Because IFNγ derived from activated CD4^pos^ T helper type 1 (Th1) and CD8^pos^ cytotoxic T cells^42^ induces and enhances CD38 expression in CD38^neg^ and CD38^pos^ AML blasts, respectively, we elected to harness the IFNγ/CD38 axis in both of these cell populations by developing a novel compact CD38-CD3 T cell engager (BN-CD38). Our data show that, even in the autologous setting, BN-CD38 activated and redirected T cells to kill CD38^pos^ AML cells, and in doing so, T cells released IFNγ directly in the tumor site, which resulted in the re-expression and upregulation of their therapeutic target, even in AML cells that initially lacked CD38 expression. Although CyTOF immunophenotyping in both newly diagnosed and relapsing patients showed that BN-CD38 significantly reduced AML blasts by concomitantly expanding autologous CD4^pos^ and CD8^pos^ T cells, BN-CD38 did not cause the reduction of CD38^pos^ non-cancer cell populations, including NK, NKT, or B cells, an effect likely due the absence of CD38 upregulation upon IFNγ treatment in these populations (data not shown).

Interestingly, several studies have investigated the preclinical activity of anti-CD38 monoclonal antibodies, including the FDA-approved antibody daratumumab, in AML as single agents and in combination with ATRA or cytarabine (a chemotherapeutic agent), but results have been contradictory.^27,43–46^ Whereas some studies have shown that targeting CD38 on AML cells using daratumumab or daratumumab/ATRA had a robust anti-leukemic activity,^45,46^ daratumumab/ATRA/cytarabine yielded no observable benefits in AML patient derived xenograft models (PDX).^44^

CD38-directed CAR T cells have also been developed and tested as a potential AML therapy. It was shown in pre-clinical setting that CD38-CAR T cells mediated killing against CD38^pos^ AML cancer cells, but CD38^neg^ AML cells escaped targeting ^47^. To mediate killing of CD38^neg^ AML cells, a combination of anti-CD38 CAR T-cell therapy with ATRA was necessary to confer complete remissions.^47^

Our work showed that IFNγ treatment of AML primary cells also upregulated vital metabolic pathways involved in LSC survival such as oxidative phosphorylation and fatty acid metabolism, concomitantly eliciting anti- and pro-survival pathways, as also previously reported ^48,49^ This finding suggests that IFNγ monotherapy may not be successful in all patients and the use of CD38 T cell engagers may be necessary to eradicate leukemia. BN-CD38 markedly suppressed AML cell engraftment and completely eliminated leukemia in mice xenografted with both AML cell lines and primary cells independently of the TP53 status of the blasts, further supporting the translational significance of these findings.

To the best of our knowledge, herein we show for the first time that IFNγ induced the upregulation of CD38 in AML blasts, including LSC-enriched CD34^pos^CD38^neg^ cells, but not in normal HSCs. To this end, we show that a novel single-chain BN-CD38 that activated both CD8- and CD4-positive T cells against CD38^pos^ AML blasts, induced release of IFNγ, thereby converting LSC-enriched CD34^pos^CD38^neg^ cells into CD34^pos^CD38^pos^ cells. These cells, despite the initial absence of the CD38 target, in turn became CD38-positive and were eliminated by the BN-CD38 engager. The net result was a robust antileukemic effect and a significant depletion of LSCs enriched in the CD34^pos^CD38^neg^ fraction. Importantly, this result did not lead to significant depletion of CD34^pos^CD38^neg^ HSCs, which we showed to be less sensitive to IFNγ, and therefore avoid killing by BN-CD38 killing. Our findings provide the scientific rationale for rapidly translating CD38 T cell engagers into the clinic as an effective antileukemic approach to deplete AML LSCs.

## Methods

### Single cell RNA-seq library preparation and bioinformatics analysis

AML and HD BM MNCs were thawed and suspended in 20% FBS IMDM medium. One million MNCs were plated in 24-well plate and total of two wells were plated for HD and AML patient. IFNγ was added to one of the wells at 10 ng/ml for 5 hours. At 5 hours, cells were collected and processed using the 10x Genomics Chromium Controller and the Chromium Single Cell 3’ Library & Gel Bead Kit (PN-120237) following standard manufacturer’s protocol. Final library quality was assessed using an Agilent Tapestation. Samples were then sequenced on the Illumina NovaSeq S4 with a read length of 2□×□50 paired end reads. Single cell RNA-seq data were processed by the CLC Bio software package. Specifically, AML and HD untreated and IFNγ treated were normalized together into one dataset matrix, predicting cell types with a CLC Bio classifier, reducing the dimensionality and clustering by UMAP, and associating cell type clusters with UMAP. Differential expression analysis was performed in the cluster of cancer cells between the treated and untreated AML samples (small number of cells in that cluster from the healthy samples were excluded). Differentially expressed genes were filtered such that the minimal number of cells whereby the expression occurred was 300 and Bonferroni corrected p-value was equal or less than 0.05. Differentially expressed genes were submitted to the IPA pathway enrichment analysis. To generate 3D UMPAPs, healthy and AML samples were normalized and clustered separately to investigate the cluster structure as a function of IFNγ treatment.

### BN-CD38 and BN-CD38Mut conceptualization, production, and characterization

Because of recent reports that matching the distance between the membranes of the tumor and T cell upon forming an immunological synapse produces the strongest T cell response^50^, we sought to create a highly compact T cell engager. Based on single chain Fv and Fabs, where a highly flexible peptide linker connects the light and heavy chains, we hypothesized that we could replace much of the flexible linker with an antigen-targeting domain and in doing so create a single chain CD38-CD3 T cell engager. To this end, we placed a nanobody bound to CD38 (5F1K.pdb) adjacent to the Fab (4IOI.pdb) using PyMOL to estimate the distance between the N and C termini of each component. These estimates suggested a linker length of 6 to 8 residues between each termini Fab termini and the nanobody would be sufficiently short but avoid steric strain. The molecule was built (peptide linker) and energy minimized using Schrodinger’s Bioluminate. To gauge how this biologic would engage T cells and CD38 on tumor cells, the CD38-nanobody complex was superimposed on the biologic, and the CDR loops of the Fab were placed adjacent to the CD3.

The gene encoding BN-CD38 was synthesized from IDT (Integrated DNA Technologies) and cloned into a mammalian expression vector using NEBuilder HiFi DNA Assembly (New England Biolabs). The mutation for generating BN-CD38Mut was introduced to BN-CD38 with Q5 Site-Directed Mutagenesis kit (New England Biolabs). Both were produced transiently in ExpiCHO cells (ThermoFisher Scientific). In brief, expression vectors were transfected into ExpiCHO cells, and the culture supernatant was collected after 7 days of incubation. Purification was done through a KanCapG column (Kaneka Corporation) followed by size exclusion chromatography (Cytiva Lifesciences). The purity and size of BN-CD38s were confirmed on SDS-PAGE. Surface plasmon resonance was performed on a Biacore T200 instrument (Cytiva Lifesciences) at 25°C to verify and measure the interaction between BN-CD38 and each target. CD38 or CD3 (AcroBiosystems) was immobilized on a CM5 chip (Cytiva Lifesciences) through EDC/NHS chemistry. Different concentrations of BN-CD38 were flowed over the chip in HBS-EP+ running buffer (Cytiva Lifesciences). The kinetic parameters were determined using Biacore Evaluation Software, version 3.0 (Cytiva Lifesciences). The cell binding experiment was done with the human T lymphocyte cell Jurkat and anti-human IgG antibody-Ax488 (ThermoFisher Scientific). Ax488 signal changes upon binding were detected using a flow cytometer (BD Accuri C6, BD Biosciences), and data were analyzed with FlowJo software (FlowJo LLC).

For further methods, please refer to the Supplemental Information.

## Acknowledgments

We are grateful to Tinisha McDonald, Kelly Synold, and Elena Pulkinen for technical support in this work. We also thank Hyeran Choi for assisting with FACS analysis of BIONICs constructs. We would also like to acknowledge the Drug Discovery and Structural Biology Core at City of Hope. We thank Yuriy Shostak and Christoph Pittius for the scientific and administrative support in this work.

Research reported in this publication was in part supported by the generous contributions of the Nason-Hollingsworth Project for Multiple Myeloma project (AK, FP). This research was also supported by the City of Hope Preclinical Drug Development Venture Program Award (FP, JW, GM) and partially supported by the National Institutes of Health grants R01-CA238429 (FP), R01-CA194742 (FP), R01-CA201382 (FP), and R50-CA252135 (EC).

Research reported in this publication included work performed at the City of Hope Liquid Tissue Bank, Analytical Cytometry, and Small Imaging cores supported by the National Institutes of Health under award number P30CA033572. The content is solely the responsibility of the authors and does not represent the official views of the National Cancer Institute or the National Institutes of Health.

