## Supplemental Information for "Leveraging IFNγ/CD38 regulation to unmask and target leukemia stem cells in acute myelogenous leukemia"

**Materials and Methods**

**Primary patient samples**

AML patient frozen bone marrow (BM) and peripheral blood (PB) mononuclear cells (MNCs), AML patient fresh blood samples, and healthy donor (HD) frozen MNCs from BM were obtained from the City of Hope (COH) Hematopoietic Tissue Biorepository (HTB). All patient characteristics are summarized in **Supplementary Table S1**. All subjects were enrolled at COH under institutional review board-approved protocols. Written informed consent was received from all participants prior to inclusion in the study. Healthy donor peripheral blood mononuclear cells (PBMCs) for extraction of purified T cells were obtained from a leukocyte filter collected from healthy platelet donors at the Michael Amini Transfusion Medical Center. MNCs from fresh blood of AML patients or HD leukocyte filters were isolated using Ficoll-Paque Plus (GE, Healthcare, Life Science) following the manufacturer’s instructions.

**Cell lines**

The AML cell lines THP1, SKM1, and U937 were generously provided by Dr. Ling Li’s laboratory at COH. MV-4;11 was purchased from ATCC. The T-ALL cell line MOLT-4 was generously provided by Dr. Srividya Swaminathan’s laboratory at COH. HL60 wildtype (WT) and HL60 CD38 knockout (KO) cell lines were generously provided by Dr. Andrew Yen’s laboratory at Cornell University as part of their published work ^51^. All indicated cell lines and multiple myeloma cell lines (MM.1S GFP(+)Luc(+), and JJN3) were cultured in RPMI‑1640 medium (Gibco) supplemented with 10% fetal bovine serum (FBS) (Catalog # 019K8420, Sigma) and 1% penicillin‑streptomycin (Gibco) at 37°C in 5% CO_2_. MV-4;11 was cultured in IMDM (Gibco) supplemented with 10% FBS and 1% penicillin‑streptomycin.

**GFP and luciferase lentivirus transduction of cell lines**

Lentivirus pseudotyped particles were produced by Lipofectamine 2000 (Life Technologies) mediated transfection of 293T cells with the lentivirus vector MSCV-Luciferase-EF1α-copGFP-T2A-Puro (System biosciences, SBI), the psPAX2 packaging construct, and a plasmid carrying G-glycoprotein of vesicular stomatitis virus (VSV-G) for 8 hours. After 8 hours, 293T transfected cells were washed and incubated in complete medium for 48 hours to collect viral supernatants. At 48 hours post-transfection, viral supernatants were collected and filtered through a 0.45 μm membrane. THP1, U937, SKM1, HL60WT, and HL60 CD38 KO cell lines were incubated with viral supernatant and 8 μg/mL of hexadimethrine bromide (Sigma-Aldrich) for 8 hours. After 8 hours, cells were washed and incubated at 37°C in 5% CO_2_ for 48 hours and selected with puromycin for 72 hours, then FACS-sorted into single cells; the clone that was expressing bright GFP was selected, sub-cultured, and tested for luciferase expression.

**Flow cytometry surface staining**

For cell surface expression analysis of AML or HD PB and BM MNCs, cells were washed with 1X PBS and stained for 30 minutes in ice cold FACS buffer (PBS+2%FBS) using the antibodies listed in **Supplementary Table S7**. After 30 min, cells were washed and analyzed on LSRII (Becton Dickinson) or BD LSR Fortessa X-20 (Becton Dickinson). Analyses were conducted using FlowJo™ Software (version 10.7.1) to determine median of fluorescence intensity (MFI) and percentage of different AML and T cell populations.

**Bulk RNA-seq sample preparation and bioinformatic analysis**

CD34(+) cells were isolated from BM MNCs from 3 AML patients and 3 HD using CD34 MicroBead Kit (Miltenyi Biotec, Catalog # 130-046-702) and treated with IFNγ at a concentration of 10ng/ml for overnight. Next day, cells were collected and cell pellet was resuspended in TRIZOL reagent (Cat. #15596018 Invitrogen Corporation) and total RNA was extracted and cleaned with NORGEN Biotek RNA-clean up and concentration kit. RNA quality and quantity were estimated using BioAnalyser Systems (Agilent Technologies). Samples with a RIN >8.0 were included. RNA sequencing libraries were prepared with Kapa RNA mRNA HyperPrep kit (Kapa Biosystems, Cat KR1352) according to the manufacturer's protocol. The final libraries were validated with the Agilent Bioanalyzer DNA High Sensitivity Kit and quantified with Qubit.  Sequencing was performed on Illumina HiSeq 2500 with the single read mode of 51cycle, and about 50 million reads were provided to each sample.  Real-time analysis (RTA) 2.2.38 software was used to process the image analysis.

For RNA-seq data analysis, sequence reads passing quality check were preprocessed for adaptor and polyA removal using trimmomatic v.0.39 ^52^ and fastp v.0.23.2 ^53^, followed by mapping to human reference genome hg38 with an index build containing the transcript/gene annotations using STAR v.2.7.9a ^54^. Strand-specific read counts of each gene annotated in RefGene were calculated by featureCounts function in the Subread package v.1.6.4 ^55^. To promote the accuracy of the gene expression estimates, genes with counts per million (CPM) > 1 in at least three samples were required for downstream analysis. Bioconductor package edgeR v.3.32.1 ^56^ was applied with a design matrix for testing differential expression between IFNγ-treated and untreated sample pairs based on quasi-likelihood F-tests. Statistical *p*-values were adjusted by Benjamini and Hochberg method. Genes below false discovery rate of 0.05 were considered significant. Preranked Gene Set Enrichment Analysis ^57^ was conducted for functional enrichment analysis with gene sets obtained from MSigDB v7.5.1 ^58^. Genes from edgeR results were ranked by the sign of logFC in combination with -log10 of the *p*-value for this analysis.

**BN-CD38 T cell killing assay with cancer cell lines (IC_50_ curves)**

To calculate IC_50_ curves of BN-CD38 or BN-CD38Mut and to determine the T cell killing activity of BN-CD38, T cells were purified using a Pan T cell isolation kit (Miltenyi, catalog # 130-096-535) from HD PBMCs. CD38^pos^ cancer cell lines (i.e., THP‑1 and MM.1S) were stably expressing GFP, whereas CD38^pos^ MV-4;11, MOLT-4, and JJN3 were labelled with CFSE according to the manufacturer’s recommendations. CD38^neg^ SKM1 cells were stably expressing GFP. HL60 and HL60 CD38 CRISPR knockout cell lines were stably expressing GFP and were induced to express CD38 using 1 µM all trans retinoic acid (ATRA) for 48 hours. All indicated cell lines were co-cultured with T cells and serially diluted control human IgG or BIONICS (i.e., BN-CD38 or BN-CD38Mut) (0.001 ng/ml to 100 ng/ml) overnight at 37 C at an E:T ratio of 1:1 (E: effectors [T cells]; T, target [cancer cells]). The next day, cells were washed with 1X PBS and stained with 7-AAD (BioLegend) for 10 minutes in the dark at room temperature. Additionally, minimum (T cells + cancer cells; 0% killing) and maximum (T cells + cancer cells + 50 µl Tween-20; 100% killing) controls were included to set 7AAD gates. Flow cytometry was performed on a BD LSR Fortessa X-20 (BD Biosciences) detecting GFP and 7-AAD. To calculate cancer cell percent killing, total cells were gated for GFP^pos^, and the percent of 7-AAD for each sample was determined. IC_50_ curves were generated with Graphpad prism transformation to log(ng/ml BN-CD38 or IgG concentration), normalization, and plotting with nonlinear fit of normalization (log(inhibitor) vs. response variable slope (four parameters) function).

**BN-CD38 T cell activation assay with cell lines**

To determine T cell activation with control human IgG, BN-CD38 or BN-CD38Mut, purified pan T cells from HD PBMCs were co-cultured with cancer cell lines at E:T 1:1 overnight in the presence of increasing doses of control human IgG, BN-CD38, and BN-CD38Mut (0.001 ng/ml-100 ng/ml). The next day, cocultured T cells and cancer cells were collected and subjected to surface staining for CD4, CD8, CD69, and CD25 (See **Supplementary Table S7** for antibody information.), and GFP/CFSE was used to exclude cancer cells.

**BN-CD38 T cell killing and activation assay with AML primary MNCs and autologous T cells**

Frozen MNCs from AML BM or PB were thawed in customized thawing medium (20% FBS, 1mg/ml DNAse, and 20,000 U heparin in IMDM) for 2 hours at room temperature to obtain single cells. AML MNCs were washed and suspended in IMDM supplemented with 20% FBS, 1% penicillin‑streptomycin, 200 pg/mL granulocyte-macrophage colony-stimulating factor (GM-CSF, R&D Catalog # 7954-GM-010/CF), 1 ng/mL granulocyte colony stimulating factor (G-CSF, R&D Catalog # 214-CS-005/CF), 200 pg/mL stem cell factor (SCF, Stem cell technologies Catalog # 78062.1), 1 ng/mL interleukin-6 (IL-6, Stem cell technologies Catalog # 78050.1), 25 ng/ml interleukin-3 (IL-3, Stem cell technologies Catalog # 78146 ), and 100 ng/ml Flt-3 ligand (Stem cell technologies, Catalog # 78137). Total AML MNCs were plated at 150,000-200,000 cells/well in 96-well plates and treated with 100 ng/ml of control human IgG and 0.1, 1.0, and 100 ng/ml BN-CD38 for 5 days. At 5 days, cells were harvested and subjected to AML cell surface marker staining (CD45, CD33, CD34, and CD38) and a T cell activation immunophenotyping panel (CD4, CD8, CD69, and CD25). To calculate percent killing, we performed the following calculation: [(MFI CD34 or CD33 BN-CD38active)/(MFI CD34 or CD33 control human IgG)]*100.

**Cytokine luminex assay**

THP1, HL60WT, and HL60 CD38 KO GFP^pos^ cells were co-cultured with purified pan T cells from HDs at E:T 1:1 overnight in the presence of 1.0 ng/ml of control human IgG, BN-CD38, and BN-CD38Mut. As a control, HD-derived T cells without co-culture with cancer cells were treated with 1.0 ng/ml of control human IgG, BN-CD38, and BN-CD38Mut. The next day, the supernatants were collected from co-cultures (cancer cells + T cells) and T cells alone for each treatment group. The collected supernatants were subjected to Human 10-plex cytokine immunoassay (R&D Systems, Catalog #LXSAHM-10, Lot #L140009). Similarly, total AML patient MNCs were treated with 1.0 ng/ml of control human IgG, BN-CD38, and BN-CD38Mut for 5 days, and supernatants were collected and subjected to human 10-plex cytokine immunoassay.

**Mass cytometry (CyTOF) staining and acquisition**

BM or PB MNCs from AML patients were thawed and suspended in the media described earlier (20% FBS IMDM+1% penicillin‑streptomycin+ cytokines and growth factors). AML MNCs were plated at 5-6 million cells per well in 6-well plates and treated with 1.0 ng/ml of control human IgG, BN-CD38, BN-CD38Mut, or a combination of 1.0 ng/ml BN-CD38 and 2 μg/ml αIFNγ neutralizing antibody (Mabtech, Clone # MT111W, Catalog # 3420-1N-500) and incubated for 5 days at 37°C. At 72 hours, 1.0 ml of indicated medium was added. At day 5, cells were harvested, washed, counted, and subjected to immunostaining with customized 36-surface marker metal conjugated antibodies (**Supplementary Table S5**) according to Fluidigm's CyTOF protocols for Cell-ID Cisplatin (PRD018 version 5) (Cat. 201064). Non-commercial metal-conjugated antibodies were purchased purified from Biolegend, and in-house antibodies were conjugated according to Fluidigm's protocol for Maxpar Antibody Labeling (PRD002 Rev 12). Samples were acquired, exported as FCS files, and normalized on Fluidigm's Helios (Software 7.0.5189).

**CyTOF Analysis**

Custom panel FCS files were manually cleaned up using FlowJo™ Software (Windows edition, Version 10.6. Becton Dickinson Company; 2019) and analyzed using the Cytobank© platform (https://www.cytobank.org) (Cytobank, Inc., Mountain View, CA) for gating and using tSNE plots, FlowSOM, and CITRUS analyses for clustering AML and immune cells populations. The gating strategy is shown in **Supplementary** **Table S6**.

**IFNγ intracellular detection**

THP1-GFP cells were co-cultured with HD T cells and treated with IgG, BN-CD38, and BN-CD38Mut at a concentration of 1.0 ng/ml and incubated overnight at 37°C. The next day, the cells were further incubated for an additional 6 hours with BD Golgi Stop (BD Biosciences). Phytohemagglutinin (PHA; 250 ng/ml; Sigma) and omitting BIONICs were used as positive and negative controls, respectively. At the end of the incubation, the cells were transferred to the FACS tubes, washed twice with FACS buffer, and then stained with CD3-AF700, CD4-BV711, CD8-PE-Cy7, CD45-BUV395, CD45RA-PerCP-Cy5.5, CCR7-BV786 and CD25-V450 for 20 minutes in the dark at 4°C. After surface staining, cells were washed with FACS buffer and permeabilized for 20 minutes at 4°C using a Cytofix/Cytoperm kit (BD Biosciences) and then washed twice with Perm/Wash buffer (BD Bioscience). Next, anti-human IFNγ-APC antibody was added to the cells, and the mixture was incubated for 20 minutes in the dark at 4°C. The cells were washed with Perm/Wash buffer and subsequently fixed in PBS containing 2% paraformaldehyde (BD Biosciences). For each sample, 50,000 total events were acquired on the BD LSR Fortessa X-20. Analyses were conducted using FlowJo™ Software (version 10.7.1).

**Methylcellulose colony-forming assay**

BM or PB MNCs from AML patients or HD BM MNCs were thawed and suspended in 20% FBS + 1% penicillin‑streptomycin IMDM with no cytokines or growth factors. AML or HD MNCs were plated at 0.25-0.5 million cells per well in 12-well plates and treated overnight with 1.0 ng/ml control human IgG, BN-CD38, BN-CD38Mut, or 10 ng/ml IFNγ (R&D, catalog # 285-IF). The next day, cells were harvested and suspended in 2% FBS + 1% penicillin‑streptomycin IMDM at a concentration of 50000 cells/200 μl, and 1.3 ml of Methocult H4034 (Stem Cell Technologies) was added. This mixture was equally divided into two wells (750 μl/ well, duplicate wells per treatment). Colonies were analyzed after 12-14 days using Widefield Zeiss Observer 7 inverted microscope at 2.5X and 5X magnification.

**Reverse transcription and quantitative real time PCR (qRT-PCR)**

Total cell pellets were suspended in TRIZOL reagent (Cat. #15596018 Invitrogen Corporation), and RNA extraction was performed using NORGEN Biotek RNA-clean up and concentration kit (catalog # 43200). cDNA synthesis was performed using the High-Capacity cDNA Reverse Transcription Kit (Applied Biosystems, Cat# 4368814) in Mastercycler pro. Quantitative real time-PCR (qRT-PCR) was performed using TaqMan Fast Advanced Master Mix (Applied Biosystems), according to the manufacturer’s instructions. The appropriate TaqMan probes for mRNA quantification were purchased from Applied Biosystems, including GAPDH used as endogenous control (HS99999905_m1) and CD38 (Hs01120071_m1), and all reactions were performed in triplicates.

***In vivo* studies**

All animal protocols were approved by the Animal Care and Use Committee of the COH, in accordance with the National Institute of Health Guidelines for the Care and Use of Laboratory Animals.

**CD38 knockout mouse model transduced with AML onco-fusion gene MLL-AF9**

Both wild-type (WT; C57BL6) and CD38-knockout mice (CD38KO Jax #003727, B6.129P2-Cd38tm1Lnd/J (homozygous)) were obtained from Jackson Laboratory. Female, 8 week old mice were used for experiments. Bone marrow LSK (Lin(-)c-Kit(+)Sca-1(+)) cells were purified from BM MNCs of WT and CD38 KO mice and transduced with retroviral vectors co-expressing MLL-AF9 (MA9) plus EGFP. GFP-positive cells were sorted at 72 hours, and a colony forming cell (CFC) assay was performed. After one round of CFC assay, the cells were transplanted into irradiated (650 cGy) B6.Ly5.1+ recipients (2X10^5^/mouse) by intravenous (IV) route. We monitored the engraftment ratio (mouse CD45.2) in peripheral blood, and once the engraftment ratio reached 10% or the mice showed any signs of illness, we harvested all the mice, and engraftment of donor cells (mouse CD45.2) in bone marrow and spleen was analyzed through flow cytometry. The spleens were collected and weighed.

**AML cell lines THP1 and U937 xenografts**

### Immunodeficient NOD.Cg-*Prkdc^scid^ Il2rg^tm1Wjl^*/SzJ (NSG) mice were used for AML cell line-derived xenografts. One million THP1-GFP^pos^Luc^pos^ cells were IV injected into NSG mice. On day 18, mice were subjected to bioluminescence imaging (BLI) and randomized into two groups with comparable tumor burden: control human IgG (n =10) or BN-CD38 (n=8). On day 19, the first treatment was administered. Each treatment included 2.5 mg/kg/mouse control human IgG or BN-CD38 and 3 million HD-derived purified T cells by IV. Control human IgG was administered twice, whereas BN-CD38 was administered four times. Weekly treatments were administered. Mice were subjected to BLI every week to monitor tumor burden.

### NSG mice were IV administered 0.5 million U937-GFP^pos^Luc^pos^ cells. On day 4, mice were blindly randomized into three groups: control human IgG (n=6), BN-CD38 (n=6), or BN-CD38Mut (n=5), and treatment with similar doses as with THP1 was IV injected weekly. Control human IgG and BN-CD38Mut were administered twice (days 4 and 11), whereas BN-CD38 was administered three times (days 4, 11, and 18). BLI was done every week to monitor tumor burden. Engraftment was checked in bone marrow using flow cytometry.

**AML primary patient-derived xenografts (PDX)**

For the PDX-1 (AML patient 14, Supplementary Table S1), CD34^pos^ cells from AML primary MNCs were injected into irradiated (150 cGy) NSG mice (1×10^6^ cells per mouse). We monitored the engraftment (human CD45^pos^CD33^pos^) in peripheral blood, and once the engraftment ratio reached 1% (day 20), we randomized the mice into three groups: control human IgG (n=5), BN-CD38 (n=5), and BN-CD38Mut (n=5). We administered three treatments (days 21, 28, and 35) by IV. Each treatment comprised 2.5 mg/kg/mouse control human IgG, BN-CD38, or BN-CD38Mut and 3 million healthy donor-derived T cells. On day 42, we harvested the bone marrow and spleen and assessed engraftment of AML cells (human CD45^pos^CD33^pos^) by flow cytometry. BM MNCs isolated from each treatment group were injected into secondary mice (one million BM MNCs per mouse). Mice were administered three treatments of control human IgG or BIONICs and HD T cells by IV with doseS described earlier. On day 27, BM and spleen were harvested to analyze AML cell engraftment by flow cytometry, and spleens were weighed to assess splenomegaly. BM MNCs harvested from each treatment group were IV injected into irradiated tertiary mice (one million cells per mouse). No treatment was administered on tertiary transplantation; only survival was monitored.

For the human AML PDX-2 (AML patient 7, Supplementary Table S1), CD34+ cells from AML primary specimen (PB8) were IV injected into irradiated (150 cGy) NSG mice (1×10^6^ cells per mouse). We monitored the engraftment (human CD45+CD33+) in NSG peripheral blood, and once the engraftment ratio reached 1% (day 21), we randomized the mice with comparable tumor burden into three groups: control human IgG (n=5), BN-CD38 active (n=5), and BN-CD38Mut (n=5). We administered two treatments (day 22 and day 29) with dose described earlier (IV co-injection of 2.5/kg/mouse control human IgG or BN-CD38 active or BN-CD38Mut and 3 million T cells purified from HD PBMCs). Engraftment of AML patient cells in bone marrow and spleen of NSG mice was analyzed by flow cytometry on day 32. We also weighed spleen to assess splenomegaly.

**Statistical analysis**

All statistical analyses were performed using GraphPad Prism (version 9, GraphPad Software Inc., San Diego, California, USA). The two-tailed paired or unpaired Student’s t-test was used to compare between two groups. The Mann-Whitney statistical test was used to compare between HD and AML patients. Ordinary one-way ANOVA was used to compare three groups. The log-rank (Mantel-Cox) was used to assess significant differences in mice survival between treatment groups. A *p* value less than 0.05 was considered statistically significant. All experiments were repeated at least three independent times, and data are presented as mean ± SEM.

**Supplemental Figures**

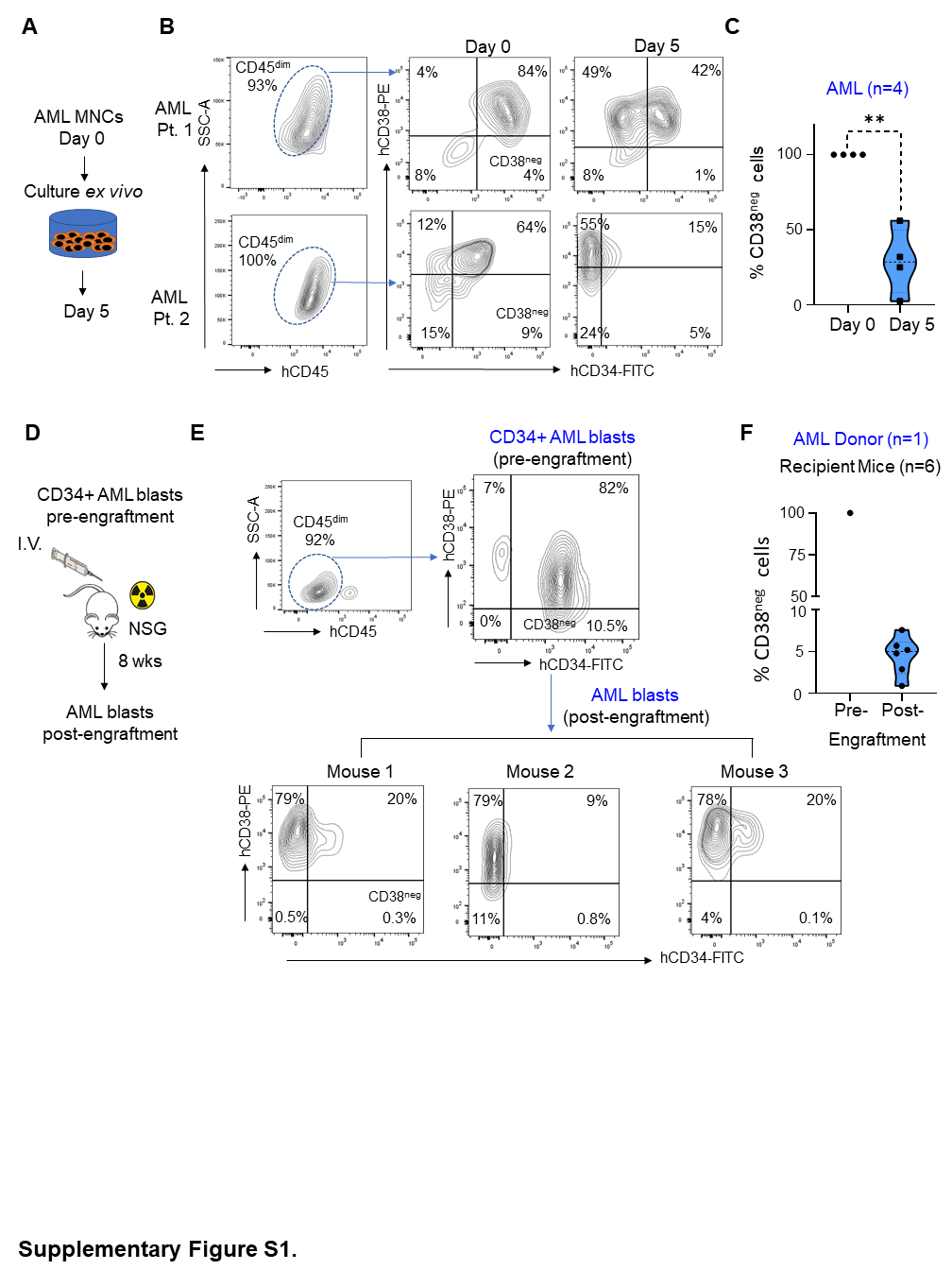

**Supplementary Figure S1.**

(A) To determine dynamics of CD38 in AML patient samples, AML primary cells were cultured *ex vivo* for 5 days (excluding one sample that was cultured for 48 hours). At day 0 and day 5, AML subpopulations were assessed by surface staining of CD34-FITC, CD38-PE, and CD45-APC. (B) Representative CD45^Dim^CD34CD38 flow cytometry contour plot of samples from two AML patients showing CD34^pos^CD38^neg^ population acquiring CD38 at day 5 of *ex vivo* culture compared to day 0. (C) Violin plot showing CD34^pos^CD38^neg^ population in four AML samples post 5 days of *ex vivo* culture acquiring CD38. (D) Schematic of assessing AML CD38 dynamics *in vivo.* AML patient cells (donor) were intravenously (I.V.) injected into irradiated NSG mice, and percent of AML human cells were assessed post-engraftment at 8 weeks in BM of NSG mice using three color hCD34-FITC, hCD38-PE, and hCD45-APC surface staining followed by flow cytometry and compared to AML patient cell immunophenotype before engraftment into mice. (E) Flow cytometry contour plot of AML sample (donor, top) before engraftment and after engraftment into NSG mice (bottom) showing CD34^pos^CD38^neg^ population acquiring CD38 post-engraftment in 3 representative mice. (F) Violin plot showing CD34^pos^CD38^neg^ population acquiring CD38 in the BM of 6 NSG mice post-engraftment compared to pre-engraftment. Data are represented as mean ± SEM.

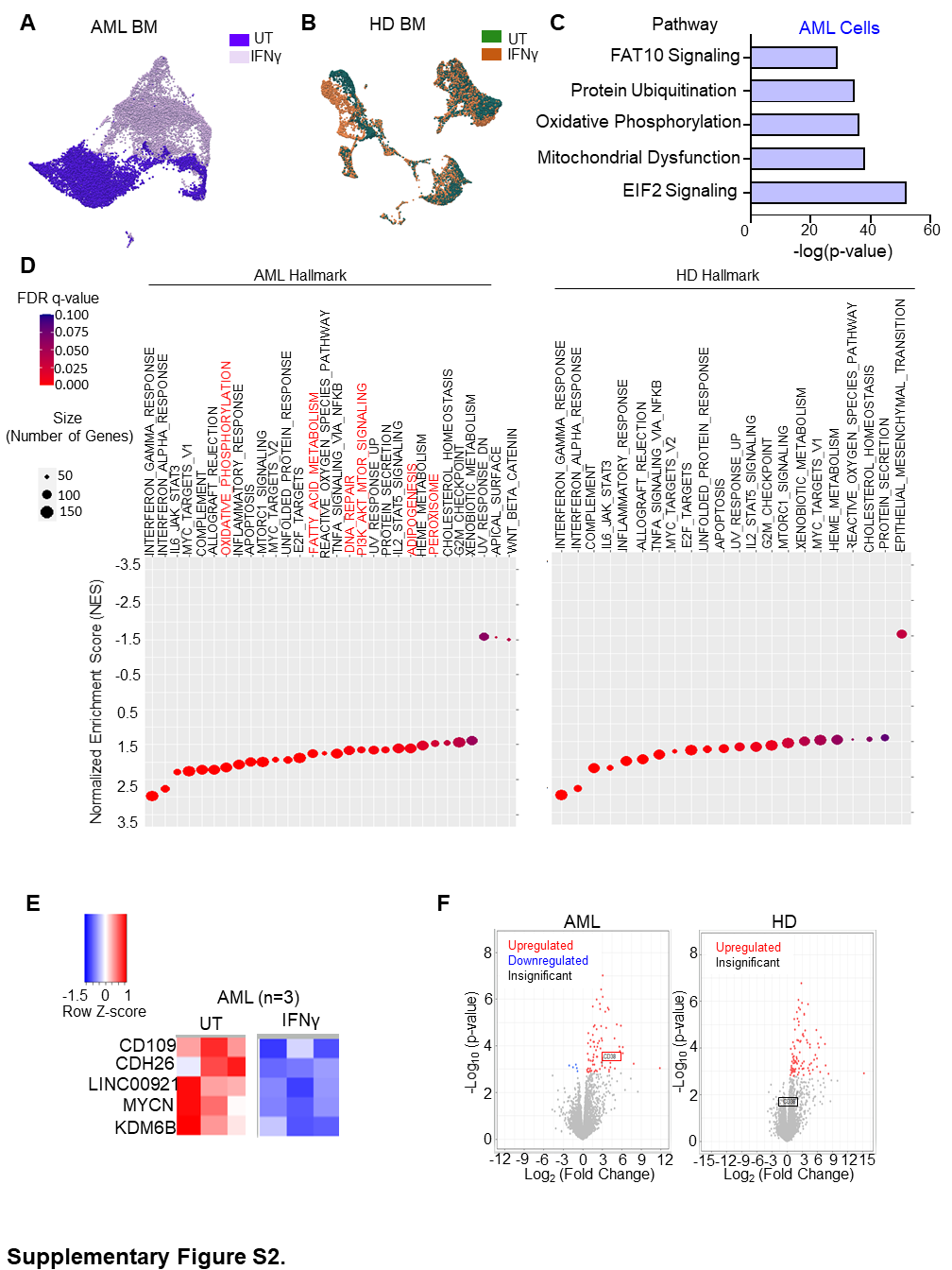

**Supplementary Figure S2.**

(A-B) 3D UMAP depicting effect of IFNγ treatment (10 ng/ml, 5 hours) on AML and HD BM MNCs. (C) Bar graph showing differentially regulated pathways in IFNγ (10 ng/ml, 5 hours) treated versus untreated AML cells in one AML sample that was treated for 5 hours with IFNγ and subjected to single cell sequencing analysis. (D-F) CD34^pos^ cells were purified from AML patients (n=3) and healthy donors (HD) (n=3) and treated overnight with IFNγ at 10 ng/ml. The next day, RNA was extracted from untreated control and IFNγ treated cells and subjected to bulk RNA-seq. (D) Normalized enrichment score (NES) plot for most positively and negatively enriched Hallmark Signature pathways identified by gene set enrichment analysis (GSEA) in HD and AML CD34^pos^ cells treated with IFNγ (FDR<0.05). (E) Heatmap of 5 downregulated genes upon IFNγ treatment in AML cells. (F) Volcano plots depicting differential gene expression upon IFNγ treatment in AML patients and HD (FDR < 0.25; p <0.05). CD38 gene expression is shown in the square for AML and HD.

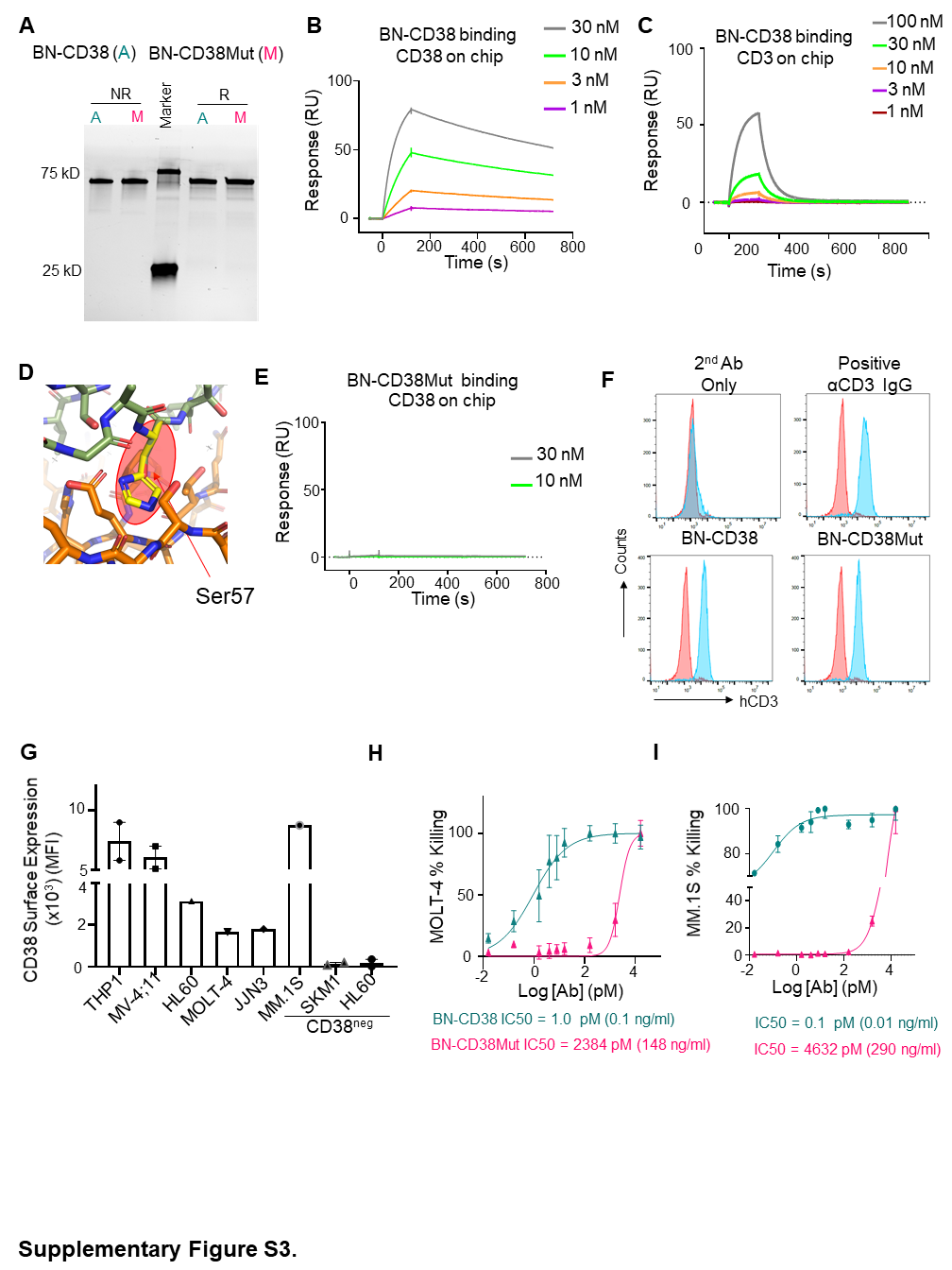

**Supplementary Figure S3**.

(A) SDS-PAGE for active (A) and mutant (M) BN-CD38. (B) SPR sensorgram for CD38 binding with different concentrations of BN-CD38. (C) SPR sensorgram for CD3 binding with different concentrations of BN-CD38. (D) Binding interface shown in licorice presentation (PDB, 5F1K), CD38 in orange, CD38 nanobody in green, and Ser57 in yellow. (E) SPR sensorgram for CD38 binding with two concentrations of BN-CD38 Mut. (F) Histogram of CD3 binding to Jurkat cells with 100 µg/mL of BN-CD38 and anti-CD3 antibody (blue) and no labeling (red). (G) CD38 surface density was assessed by flow cytometry on AML (THP1, MV-4;11, and SKM1), APL (HL60 and HL60 CD38 knockout), T-ALL (MOLT-4), and MM (MM.1S and JJN-3) cell lines. HL60 CD38WT GFP^pos^ and HL60 CD38KO GFP^pos^ cell lines were treated with 1 μM ATRA for 48 hours before determining CD38 expression. (H-I) MOLT-4 CFSE^pos^ and MM.1S GFP^pos^ cells were co-cultured with healthy donor T cells at an E:T ratio of 1:1 overnight (16hrs) in the presence of increasing doses of BN-CD38 and BN-CD38Mut. To assess MOLT-4 and MM.1S killing by effector T cells, MOLT-4 and MM.1S cell viability was determined with 7-AAD staining and gating on CFSE^pos^ or GFP^pos^ cells by flow cytometry. IC_50_ curves are shown for BN-CD38 and BN-CD38Mut, and data are represented as mean ± SEM of 3 independent healthy donors. (Target cancer cells, T; Effector T cells, E; all trans retinoic acid, ATRA; MM, multiple myeloma; APL, Acute promyelocytic leukemia).

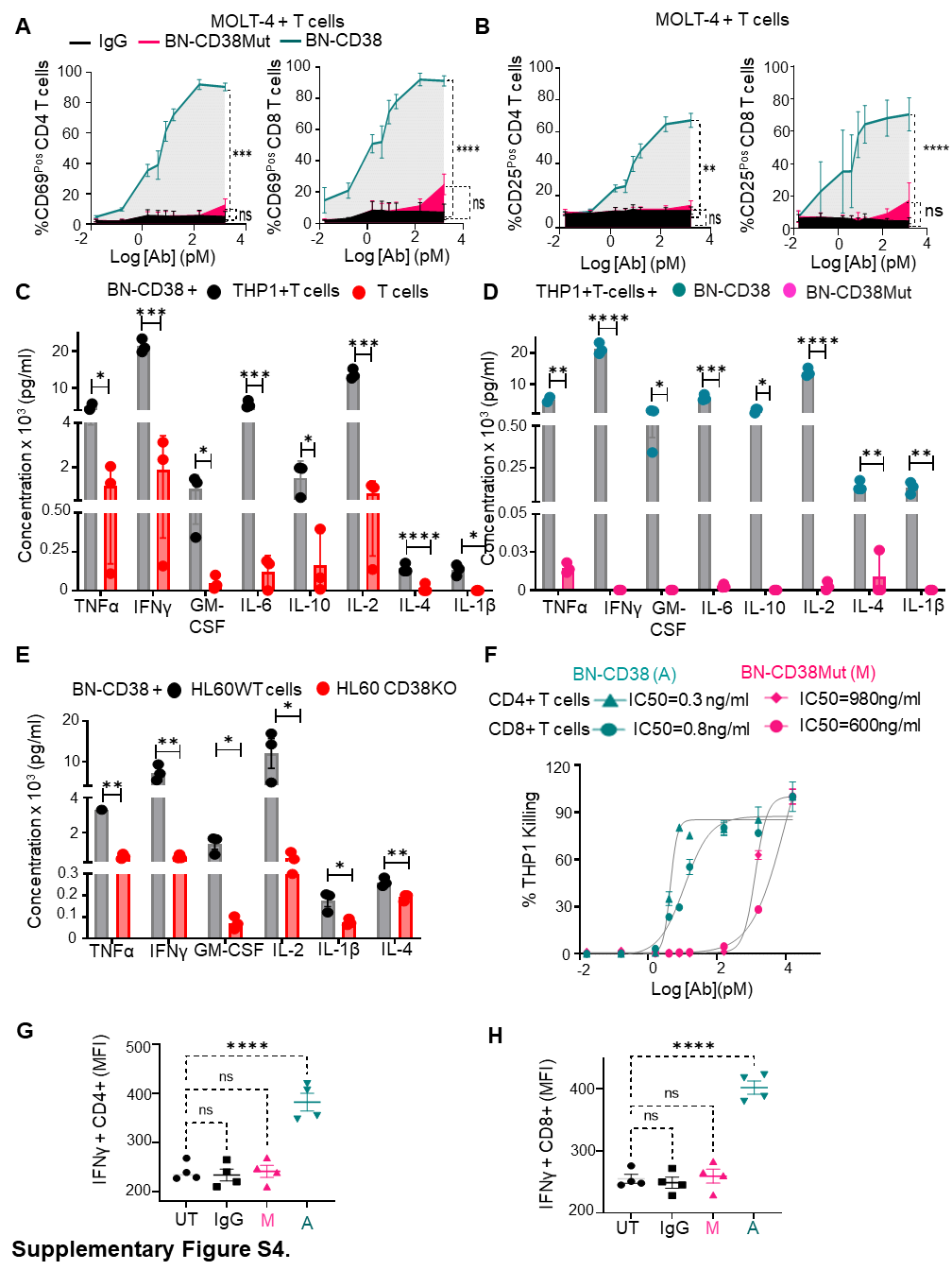

**Supplementary Figure S4**.

(A-B) MOLT-4 CFSE^pos^ cells were co-cultured with healthy donor T cells at an E:T ratio of 1:1 overnight (16hrs) in the presence of different doses of BN-CD38, BN-CD38Mut, and control human IgG. Early and late T cell activation was assessed by surface expression of CD69 and CD25 in CD4^pos^ and CD8^pos^ subsets with flow cytometry and by excluding CFSE^+^/GFP^+^ cancer cells. Dose-dependent T cell activation curves are represented as mean ± SEM of 3 independent healthy donors. Ordinary one-way ANOVA with Dunnett's multiple comparisons test was used for calculation of statistical significance. **p<0.01; ***p<0.001; ****p<0.0001; ns, not significant. (C-D) THP1 GFP^pos^ cells were co-cultured with healthy donor T cells overnight (16hrs) at an E:T ratio of 1:1 in the presence of 1.0 ng/ml of BN-CD38 and BN-CD38Mut. At 16 hours, supernatants were harvested from corresponding wells and subjected to Human 10-Plex cytokine immunoassay. Bar graphs compare concentration (pg/ml) of cytokines produced in supernatants of THP1-T cell co-cultures with BN-CD38 versus T cells alone in the presence of 1.0 ng/ml BN-CD38 (C), or versus THP1-T cell co-cultures with BN-CD38Mut (D). Data are represented as mean ± SEM from 3 healthy donors. Statistical significance was calculated with unpaired student t-test. *p<0.05; **p<0.01; ***p<0.001; ****p<0.0001. (E) HL60 GFP^pos^ and HL60 CD38 CRISPR knockout GFP^pos^ cell lines were treated with 1 μM ATRA for 48 hours and co-cultured with healthy donor T cells overnight (16hrs) at an E:T ratio of 1:1 in the presence of 1.0 ng/ml of BN-CD38. At 16 hours, supernatants were harvested from co-cultures and subjected to Human 10-Plex cytokine immunoassay. Bar graphs compare concentration (pg/ml) of cytokines produced in supernatants of HL60 GFP^pos^ versus HL60 CD38 CRISPR knockout GFP^pos^ cells co-cultured with T cells in the presence of 1.0 ng/ml BN-CD38. Statistical significance was calculated with unpaired student t-test. *p<0.05; **p<0.01. (F) THP1-GFP^pos^ cells were co-cultured with purified healthy donor CD4^pos^ and CD8^pos^ T cells at an E:T of 1:1 overnight with increasing concentrations of BN-CD38 or BN-CD38Mut. THP1 cell killing by effector CD4^pos^ or CD8^pos^ T cells was determined with 7-AAD staining and gating on GFP^pos^ cells by flow cytometry. IC_50_ curves are shown for BN-CD38 and BN-CD38Mut. (G-H) THP1-GFP^pos^ cells were co-cultured with purified healthy donor T cells overnight in the presence of 1.0 ng/ml of control human IgG, BN-CD38, and BN-CD38Mut, and intracellular IFNγ was measured in CD4^pos^ and CD8^pos^ T cells by flow cytometry. Data are represented as mean ± SEM for two representative healthy donors, and each sample was used in duplicate. Ordinary one-way ANOVA with Dunnett's multiple comparisons test was used to calculate significance. ****p<0.0001; ns, not significant. (Target cancer cells, T; Effector T cells, E; all trans retinoic acid, ATRA).

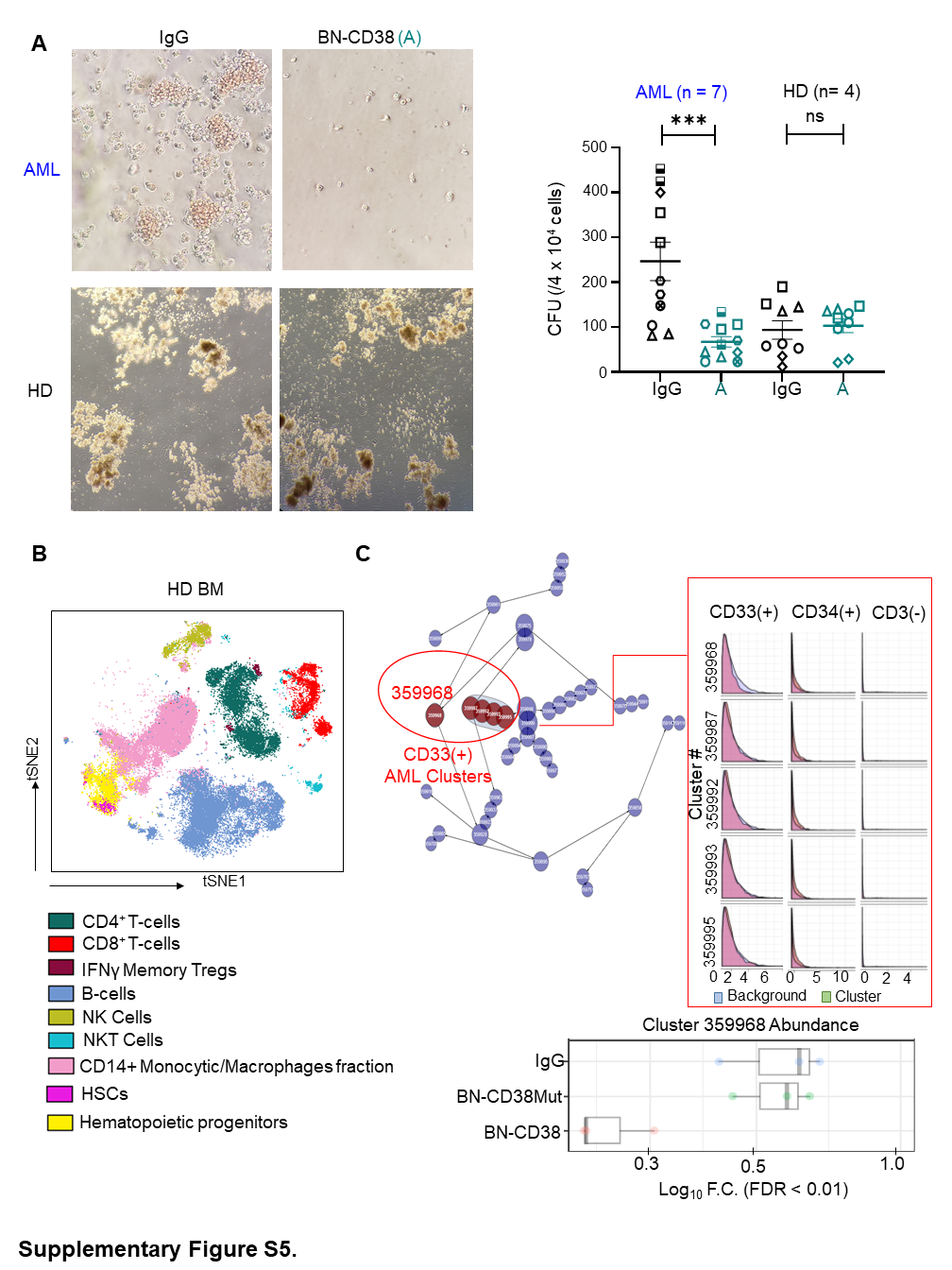

**Supplementary Figure S5**.

(A) (Left) Representative CFC assay images of AML and HD MNCs treated with 1.0 ng/ml control human IgG and BN-CD38 (A). Images were acquired at 10X magnification. (Right) Scatter plot showing CFU for AML MNCs (n= 7: 4 PB and 3 BM) and HD (n= 4 BM) MNCs treated with 1.0 ng/ml control human IgG or BN-CD38 (A). Each shape represents an AML patient sample and HD (done in duplicates). Error bars are represented as mean ± SEM. Ordinary one-way ANOVA with Dunnett's multiple comparisons test was used to calculate significance. ***p<0.001; ns, not significant. (B) Supervised high-fidelity FlowSOM (“self-organizing maps”) based on vi-SNE 2D analysis showing lymphocyte and HSC populations in HD bone marrow. (C) (Left) Unsupervised CITRUS clustering analysis of 3 CD34^pos^ AML PB samples revealed 5 clusters (red circles) were significantly changed (FDR<0.01) between BN-CD38 and control groups (IgG and BN-CD38Mut). (Right) Marker expression (histograms) of 5 red clusters showed to be enriched in AML expression markers CD33 and CD34 and negative for other immune subsets surface markers such as CD3. (Bottom) Representative bar graph of one of five red clusters showing significant (FDR < 0.01) decrease in abundance of AML cluster 359968 with BN-CD38. (MNCs= mononuclear cells; PB= peripheral blood; BM= bone marrow; CFC = Colony forming cell assay; CFU = colony forming unit).

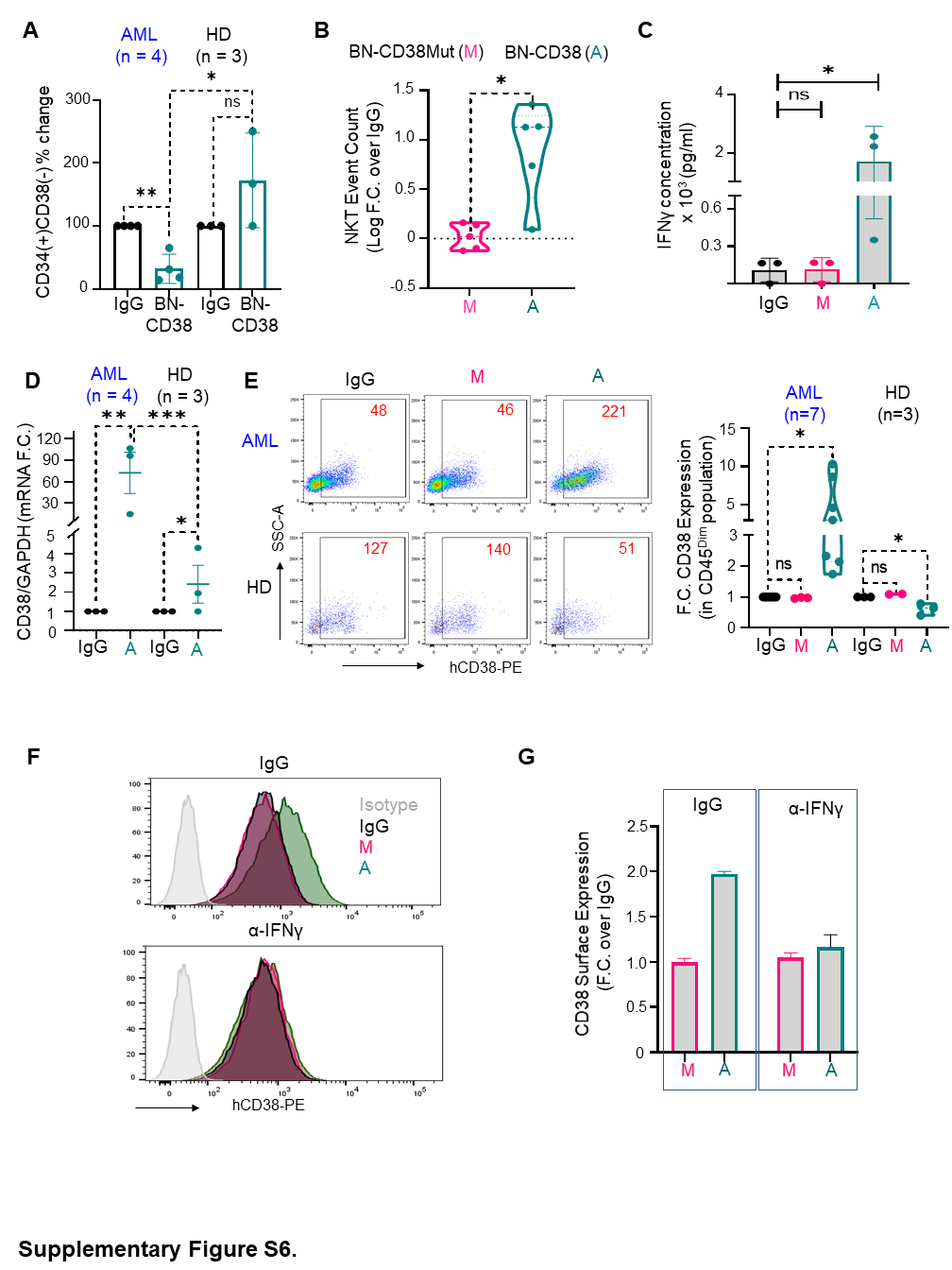

**Supplementary Figure S6**.

(A) Bar graph depicting percent of CD34^pos^CD38^neg^ population in AML (n= 4: 3 PB and 1 LP) and HD (n=3 BM) MNCs treated with control human IgG or BN-CD38 for 48 hours. Data are represented as mean ± SEM. Paired student t-test is used for calculation of statistical significance between untreated and BN-CD38 treated groups within HD and AML, whereas unpaired student t-test was used to compare AML versus HD BN-CD38 treated groups. *p<0.05; **p<0.01; ns, not significant. (B) Violin plot showing active BN-CD38, but not BN-CD38Mut, expands NKT cells. Event counts of BN-CD38 and BN-CD38Mut were normalized to control human IgG, and normalization was converted to log scale. (C) Total MNCs from three AML patients were treated with BN-CD38 (A), BN-CD38Mut (M), and control human IgG for 5 days, and supernatants were collected and subjected to Human 10-Plex cytokine immunoassay. Bar graph depicts IFNγ concentration in pg/ml for three treatment groups for three AML patients. (D) Scatter plot showing CD38 mRNA expression determined by qRT-PCR in AML (n=3: 2 PB and 1 BM) versus HD (n= 3 BM) total MNCs treated with 1.0 ng/ml control human IgG or BN-CD38 for 48 hours. Each AML patient and HD qPCR was done in triplicate; error bars represent mean ± SEM. Paired student t-test was used to compare significance between treatment groups, whereas unpaired t-test was used to calculate statistical significance between AML patients and HD. *p<0.05; **p<0.01; ***p<0.001. (E) (Left) Representative dot plot showing CD38 expression in CD45^Dim^ population following treatment of bulk MNCs with 1.0 ng/ml of BN-CD38 (A), BN-CD38Mut (M), or control human IgG. (Right) Violin plot comparing CD38 surface expression in CD45^Dim^ population of AML (n= 7: 5 PB, 1 BM, and 1 LP) and HD (n=3 BM) MNCs treated with 1.0 ng/ml of control human IgG, BN-CD38Mut (M), or BN-CD38 (A) for 48 hours. Ordinary one-way ANOVA with Dunnett's multiple comparisons test was used to calculate significance. *p<0.05. (F-G) THP1 cells were co-cultured with purified healthy donor T cells and treated with 1.0 ng/ml of BN-CD38, BN-CD38Mut, or control human IgG overnight, and supernatants were collected. Supernatant of each treatment group was added to new THP1 cells in presence of IgG and αIFNγ neutralizing antibody (2 μg/ml) for 48 hours ,and CD38 surface expression was determined by flow cytometry, which are represented by histograms and bar graph. Data are represented as mean ± SEM of two independent experiments.

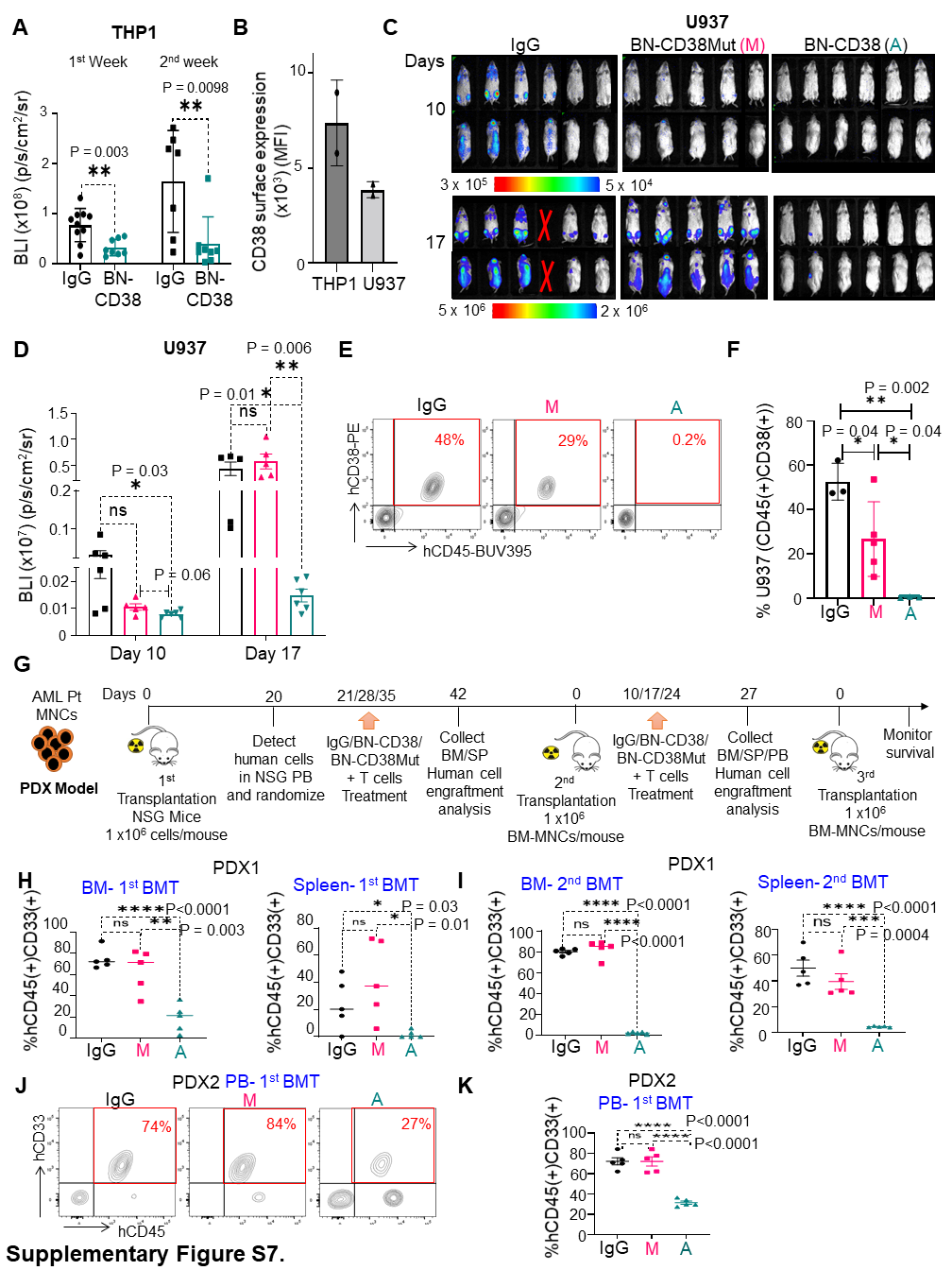

(A) Radiance values were calculated from BLI data for control human IgG and BN-CD38 treated THP1 xenografts on days 25 and 29 and are illustrated in bar graphs. Error bars are represented as mean ± SEM. Unpaired student t-test was used for statistical significance calculation. Week 1 **p=0.003; Week 2 **p=0.0098 (B) Bar graph comparing CD38 surface expression in THP1 versus U937 by MFI. (C-F) U937 cells, an aggressive AML model, stably expressing luciferase were intravenously (IV) injected into NSG mice (0.5 million cells/mouse). On day 4, mice were blindly (without using BLI) randomized into 3 groups: control human IgG (n = 6), BN-CD38 (n = 6), and BN-CD38Mut (n = 5). Treatment was started on day 4, and mice were administered 2.5 mg/kg/mouse control human IgG, BN-CD38, or BN-CD38Mut together with 3 million healthy donor derived human T cells/mouse weekly by IV, for a total of 3 treatments of BIONICs and T cells. Mice were monitored weekly by BLI for tumor burden. (C) BLI data comparing tumor burden between control human IgG, BN-CD38Mut, and BN-CD38 treated groups on days 10 and 17 post-tumor cell injection. (D) Bar graph illustration of radiance values for three treatment groups derived from BLI data. Error bars are represented as mean ± SEM. Day 10 *p=0.03; Day 17 *p=0.01 for BN-CD38 versus IgG and **p=0.006 for BN-CD38 versus BN-CD38Mut*; ns, not significant. (E-F) BM MNCs were harvested from three treatment groups and assessed for engraftment of AML human cells. Flow cytometry surface analysis of hCD45 and hCD38 was assessed. Representative contour plots of hCD45 vs hCD38 are shown for one mouse from each treatment group. Bar graphs illustrate that BN-CD38 suppressed engraftment of U937 cells as assessed by U937 cell surface markers hCD45/hCD38 expression. *p<0.05; **p<0.01. (G) Setup and timeline for treatments of AML PDX1 model (description in Figure 4 K-L legend). (H) Scatter plot depicting BN-CD38 reduced AML cells engraftment in BM and spleen on first BMT in PDX-1. (BM, ****p<0.0001 for BN-CD38 versus IgG and **p=0.003 for BN-CD38 versus BN-CD38Mut; Spleen, *p=0.03 for BN-CD38 versus IgG and *p=0.01 for BN-CD38 versus BN-CD38 Mut; ns, not significant) (I) Scatter plots comparing engraftment of AML patient cells in BM and SP of three treatment groups on secondary transplantation in PDX1. Data are represented as mean ± SEM, and unpaired student t-test was used to calculate statistical significance. (BM, BN-CD38 versus IgG ****p<0.0001 and BN-CD38 versus BN-CD38Mut ****p<0.0001; Spleen, BN-CD38 versus IgG ****p<0.0001 and BN-CD38 versus BN-CD38Mut ***p=0.0004; ns, not significant) (J) Representative contour plots of one mouse from each treatment group showing that BN-CD38 suppressed AML cell engraftment in PB on first BMT in PDX-2. (K) Scatter plot depicting BN-CD38 reduced AML cell engraftment in PB on first BMT in PDX-2. ****p<0.0001; ns, not significant. (BLI = bioluminescence; BM = bone marrow; MNCs = mononuclear cells; PDX = patient derived xenograft; SP = spleen; BMT = bone marrow transplantation; MFI median fluorescence intensity).

**Table S1.** Characteristics of AML patients used in the study.

| **Assigned ID** | **Disease Status** | **Gene Mutation** | **Cytogenetics** | **Age** | **Gender** | **% Blasts** | **Used in Figure** |
| --- | --- | --- | --- | --- | --- | --- | --- |
| 1 | Relapsed | BCOR  EP300  NRAS KMT2A-MLLT10 (Fusion)  TBL1XR1-TP63 (Fusion) | ins(10;11)(p12;q23.3q23.3),+19 | 58 | M | 95% | Fig. 1B and 1C  Fig. 2B-2E Supplementary Fig. S5C Supplementary Fig. S6C |
| 2 | Refractory | FLT3-ITD  KMT2A-TET1 (Fusion) NRAS  PTPN11 WT1 | Complex Karyotype | 71 | M | 98% | Fig. 1B and 1C |
| 3 | Relapsed/Refractory | BAALC  BCOR  CDK6   DNMT3A EGFL7  ENO1-PRDM16  (Fusion) IDH2  NRAS SRSF2 | KARYOTYPE:  Clone 1:  46,XX,der(6)t(6;8)(q13;q22)[7]  Clone 2:  48,XX,+der(1)t(1;12),t(1;12)(p31.2;p13.2),+19[4]  Constitutional Cell Line:  46,XX[9]  FISH:  Positive for ETV6 rearrangement with concurrent gain of 5'ETV6, gains of CKS1B and MYC, and loss of FOXO3  Impression:  Results of current study are consistent with relapse of acute myeloid leukemia | 45 | F | 64% | Fig. 1B and 1C  Fig. 3I-3K  Supplementary Fig. S1C Supplementary Fig. S5A Supplementary Fig. S6A and S6E |
| 4 | Relapsed | ASXL1   CBL  FLT3  PHF6  SETBP1  TET2 | KARYOTYPE:  Stemline:  46,XX,add(9)(p13),del(12)(p12), t(16;22)(q22.3;q13)[16]  Sideline:  46,sl,add(15)(q22)[4]  FISH:  Positive for losses of CDKN2A and ETV6  Impression:  Results of current study consistent with relapse of acute myeloid leukemia | 61 | F | 49% | Fig. 1B and 1C  Fig. 2E-2H  Supplementary Fig. S2A, S2C, S2D-S2F  Fig. 3I-3K  Fig. 4A-4E  Supplementary Fig. S1C  Supplementary Fig. S5A and 5C Supplementary Fig. S6A and S6E |
| 5 | Newly diagnosed | CSF3R DNMT3A  MCL1 NOTCH1 | KARYOTYPE:  46,XX[20]  Impression:  Normal karyotype | 53 | F | 12% | Fig. 1B and 1C |
| 6 | Newly Diagnosed AML, transformed from relapsed MDS | TP53 | No Mitotic Cells Available For Conventional Cytogenetics  FISH:  Low level positivity for loss of EGR1 and monosomy for chromosome 7 (3.0%)  Impression:  Results of current study consistent with persistence of acute myeloid leukemia | 62 | F | 13% | Fig. 1B and 1C |
| 7 | Refractory | KIT NRAS TP53 | Complex | 32 | M | 64% | Fig. 1B and 1C  Fig. 2A-2B, 2D, 2F-2H  Supplementary Fig. S1E-S1F  Supplementary Fig. S2D-S2F Supplementary Fig. S5A Supplementary Fig. S6D-S6E  Supplementary Fig. S7J-S7K |
| 8 | Newly diagnosed | IDH1 NPM1 | Normal karyotype | 67 | M | 61% | Fig. 1B and 1C  Fig. 3I-3K Supplementary Fig. 6E |
| 9 | Newly Relapsed | FLT3  NPM1 | KARYOTYPE:  46,XY[21]  Impression:  Normal karyotype | 58 | M | 88% | Fig. 1B and 1C  Fig. 3I-3K Supplementary Fig. S5A Supplementary Fig. S6D-S6E |
| 10 | Relapsed, Progressing on treatment  (AML evolved out of Myelofibrosis) | ASXL1 CBL  JAK2 | KARYOTYPE:  46,XY[20]  Impression:  Normal karyotype | 42 | M | 23% | Fig. 1B and 1C  Fig. 4B-4G  Supplementary Fig. S5C Supplementary Fig. S6C |
| 11 | Relapsed/Refractory | CEBPA Mutation | +21del(9)(q13q22) | 54 | F | 99% | Fig. 1B and 1C |
| 12 | Newly Diagnosed | CEBPA FLT3-ITD  WT1 | Normal karyotype | 28 | M | 56% | Fig. 1B and 1C  Supplementary Fig. S1B-S1C Supplementary Fig. S6A and S6E |
| 13 | Newly Diagnosed | FLT3-ITD  NUP98-NSD1 (Fusion) | del(9)(q13q34] | 29 | M | 86% | Fig. 1B and 1C |
| 14 | Relapsed | TET2 Pos  KRAS pos.  DNMT3A | Complex Karyotype | NA | NA | 34% | Fig. 1B and 1C  Fig. 2D Fig. 4K-4L  Supplementary Fig. S7H-S7I |
| 15 | Relapsed/Refractory | FLT3-ITD | t(3;13)(p25;q12) | 41 | M | 77.8% | Fig. 1B and 1C |
| 16 | Newly Diagnosed | CBFB-MYH11 (Fusion) KIT | inv(16)(p13.1q22.1) | 20 | M | 43.6% | Fig. 1B and 1C  Fig. 4A-4E Supplementary Fig. S6C |
| 17 | Newly Diagnosed | FLT3-ITD  NPM1  TET2 | Normal karyotype | 69 | F | 90.2% | Fig. 1B and 1C  Fig. 2B Supplementary Fig. S6D |
| 18 | Newly Diagnosed | DNMT3A IDH2 KRAS NRAS NPM1 | Normal male karyotype | 40 | M | 73% | Fig. 1B and 1C  Fig. 2A-2D  Fig. 3I-3K Supplementary Fig. S5A |
| 19 | Newly Diagnosed | FLT3* NPM1 SSBP2-CHD1 (Fusion) WT1 | Normal karyotype | 38 | F | 74% | Fig. 1B and 1C  Fig. 2B and  Fig. 3I-3K Supplementary Fig. S6E |
| 20 | Newly Diagnosed | DNMT3A NPM1 TET2 | Normal karyotype | 72 | F | promonocytes (53%) with few (2%) monoblasts | Fig. 1B and 1C |
| 21 | Newly Diagnosed | IDH2 NPM1 | Normal karyotype | 67 | M | 77% | Fig. 4A-4E |
| 22 | Progressive disease | TP53 | Complex | 63 | F | 81% | Fig. 2F-2H  Supplementary Fig. S2D-S2F |

**Table S5.** CyTOF 36-metal conjugated customized antibody panel.

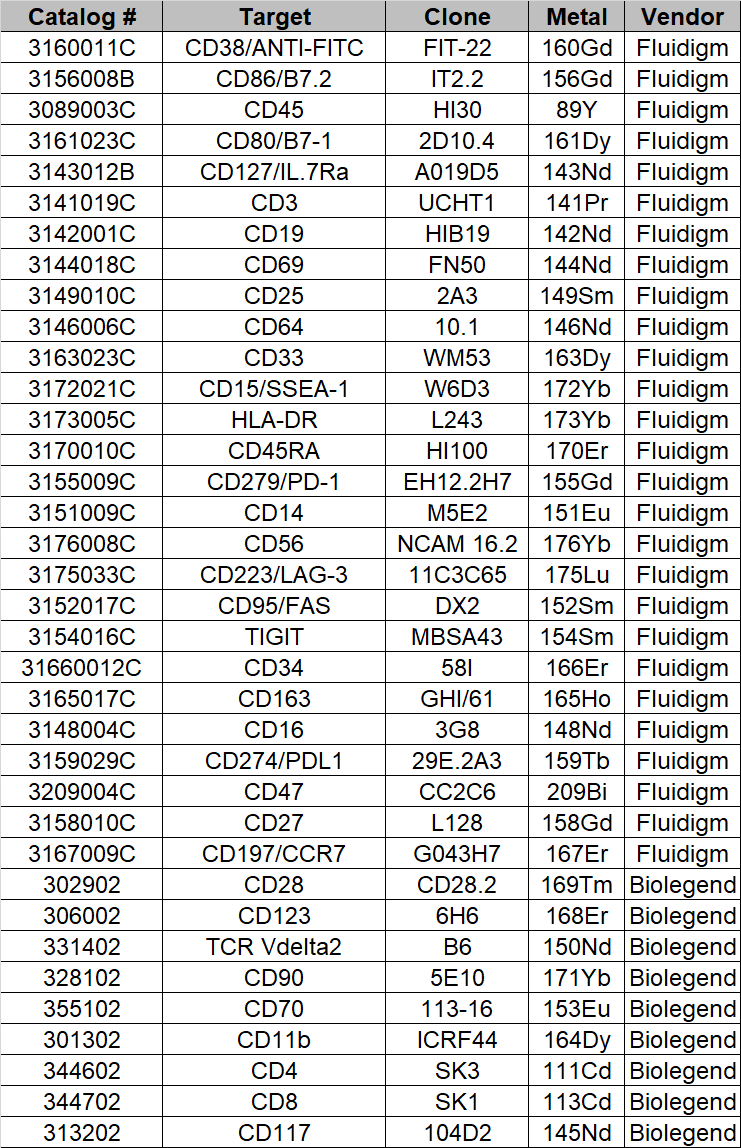

**Table S6.** CyTOF gating strategy to identify AML and immune cells subpopulations.

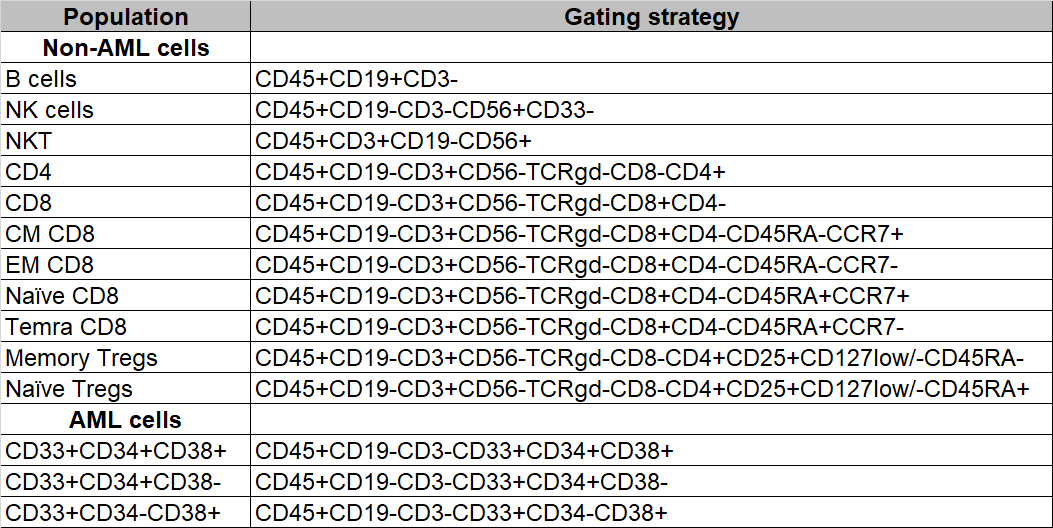

**Table S7.** Antibodies used for flow cytometry analysis.

| **Marker** | **Fluorochrome** | **Vendor** | **Clone** | **Catalog #** |
| --- | --- | --- | --- | --- |
| **Human** |  |  |  |  |
| CD45 | BUV395 | BD | HI30 | 563792 |
| CD45 | APC | BioLegend | 2D1 | 368512 |
| CD34 | APC | Invitrogen | 4H11 | 17-0349-41 |
| CD34 | FITC | BioLegend | 581 | 343504 |
| CD38 | PE | BioLegend | HB-7 | 356604 |
| CD33 | PerCP-Cy5.5 | BD | P67.6 | 341650 |
| CD4 | BV711 | BD | SK3 | 563033 |
| CD8 | PE-Cy7 | BD | RPA-T8 | 560917 |
| CD69 | APC | BioLegend | FN50 | 310910 |
| CD25 | V450 | BD | M-A251 | 560355 |
| IFNγ | APC | BD Biosciences | B27 | 554702 |
| **Mouse** |  |  |  |  |
| CD45.1 | PE-Cy7 | BioLegend | A20 | 110730 |
| CD45.2 | PE-CF594 | BD | 104 | 565390 |
